# Imaging the microscopic viscoelastic anisotropy in living cells

**DOI:** 10.1101/2023.05.28.542585

**Authors:** Hamid Keshmiri, Domagoj Cikes, Marketa Samalova, Lukas Schindler, Lisa-Marie Appel, Michal Urbanek, Ivan Yudushkin, Dea Slade, Wolfgang J. Weninger, Alexis Peaucelle, Josef Penninger, Kareem Elsayad

## Abstract

Maintaining and modulating the mechanical anisotropy is essential for biological processes. How this is achieved on the microscopic scale in living soft matter is however not always clear. Here we introduce Brillouin Light Scattering Anisotropy Microscopy (BLAM) for mapping the high-frequency viscoelastic anisotropy inside living cells. Following proof-of-principle experiments on muscle myofibers, we apply this to study two fundamental biological processes. In plant cell walls we show how a phase-transition driven switch between anisotropic-isotropic wall properties may lead to asymmetric growth. In mammalian cell nuclei we uncover a spatio-temporally oscillating elastic anisotropy correlated to chromatin condensation, with long range orientational correlations that may provide a dynamic framework for coordinating intra-nuclear processes. Our results highlight the direct and indirect role the high-frequency mechanics can play in providing dynamic structure that lead to the regulation of diverse fundamental processes in biological systems, and offer a means for studying these. BLAM should find diverse biomedical and material characterization applications.

---

In everyday life we are quite familiar with the general concept of mechanical anisotropy: a material having a different mechanical response when probed from one direction versus another. The mechanical anisotropy can often be explained if we have a certain amount of knowledge of the constituent components and their interactions. For example, those of a spring mattress may be modeled knowing the properties, abundance and orientations of its many springs and how they are connected to each other and to the mattress scaffold. This becomes more challenging in a dynamic system of mixed liquid and solid states, where the local chemistry can modulate the microscopic mechanical properties, as is the case in living cells.

Mechanical anisotropy is also essential for numerous cellular processes, such as for maintaining cell shape, motility and signaling, and diverse pathologies have been linked to changes thereof^1^. On the sub-cellular scale it can be measured by performing sequential perturbation-response measurements from different directions^2^, measuring deformations of local structures^3^, or studying asymmetries in molecular dynamics^4^. These approaches typically measure tensile or shear properties at microsecond time scales or longer, which are determined predominantly by solid constituents and scaffolds in cells.

On the microscopic scale, intermolecular interactions which have much shorter (picosecond-nanosecond) lifetimes, will ultimately determine structure in soft matter.^5–9^ Mechanics at these time scales define the supramolecular chemistry and determine the phase state of a material, the importance of which in biology is becoming increasingly apparent^10^. While for solid-like materials the mechanical anisotropy can be expected to mirror that at longer time scales, at these short time scales, fluid-like materials can also exhibit mechanical anisotropy due to solute or thermal heterogeneities. This anisotropy can in general be short-lived, on account of diffusive processes that alter the local electrostatic environment, with a persistent anisotropy suggestive of some form of structural arrest. In living cells which exist in a locally out-of-equilibrium mixed solid-fluid state, it can thus give insight into dynamic material structure and emergent phenomena on the mesoscopic scale.

Picosecond-nanosecond mechanical properties can be measured using light scattering techniques such as Brillouin Light Scattering (BLS) spectroscopy^11^. Spontaneous BLS spectroscopy is a label-free technique that can probe the high-frequency (GHz) mechanical properties by optically measuring the energies of inherent collective-molecular vibrations (acoustic phonons)^12^. These energies are obtained from the frequency shift *ν*_*B*_ of resonant acoustic phonon scattering peaks, which are typically shifted 5-15 GHz relative to a probing optical laser frequency in hydrated biological matter. *ν*_*B*_ is directly related to the isentropic elastic (*storage*) modulus, whereas the width of the Brillouin scattering peak is related to the viscous (*loss*) properties at the measured frequencies (Supplementary Text).

Being an all optical-technique, BLS has the possibility to measure the mechanical anisotropy even when direct physical access to the region of interest is not possible. This has been used to study the static structure in non-living biological materials such as collagen^13^, silk^14^, and the cornea^15^ by sequential measurements at different angles. Measuring the BLS spectrum is however non-trivial, given its close spectral proximity to elastic scattered light, and requires the use of special spectrometers that have a very high resolution and finesse^12^. While advances in Brillouin spectrometer design have opened the possibility of studying living cells^16^, current approaches are not conducive to spatio-temporal mapping of the anisotropy in living soft matter.

Here we introduce an approach for spatio-temporal mapping the GHz-frequency viscoelastic anisotropy in living cells using single-shot angle resolved Brillouin Light Scattering Spectroscopy. We apply this for studying two distinct cellular components, one where from structural considerations an anisotropy is expected (cell walls), and one where this is less clear (cell nucleus). In both cases we uncover a previously inaccessible dynamic mechanical landscape that appears to serve a fundamental biological purpose.

## Results

### Angle-resolved BLS imaging

We overcome the limitations of existing approaches for performing angle-resolved BLS by using a custom-fabricated dispersive element, based on the Virtual Imaged Phased Array (VIPA)^12, 16^, which allows one to simultaneously measure the BLS spectra from different in-plane scattering angles (Fig 1A, Methods, Supplementary Fig 1). The generated 2D radial spectral projection at each probed position gives a snapshot of the mechanical anisotropy, which can be confocally scanned through a sample to generate spatial maps with a transverse and axial spatial resolution of <2 μm (Methods, Supplementary Fig 2).

**Fig. 1.**
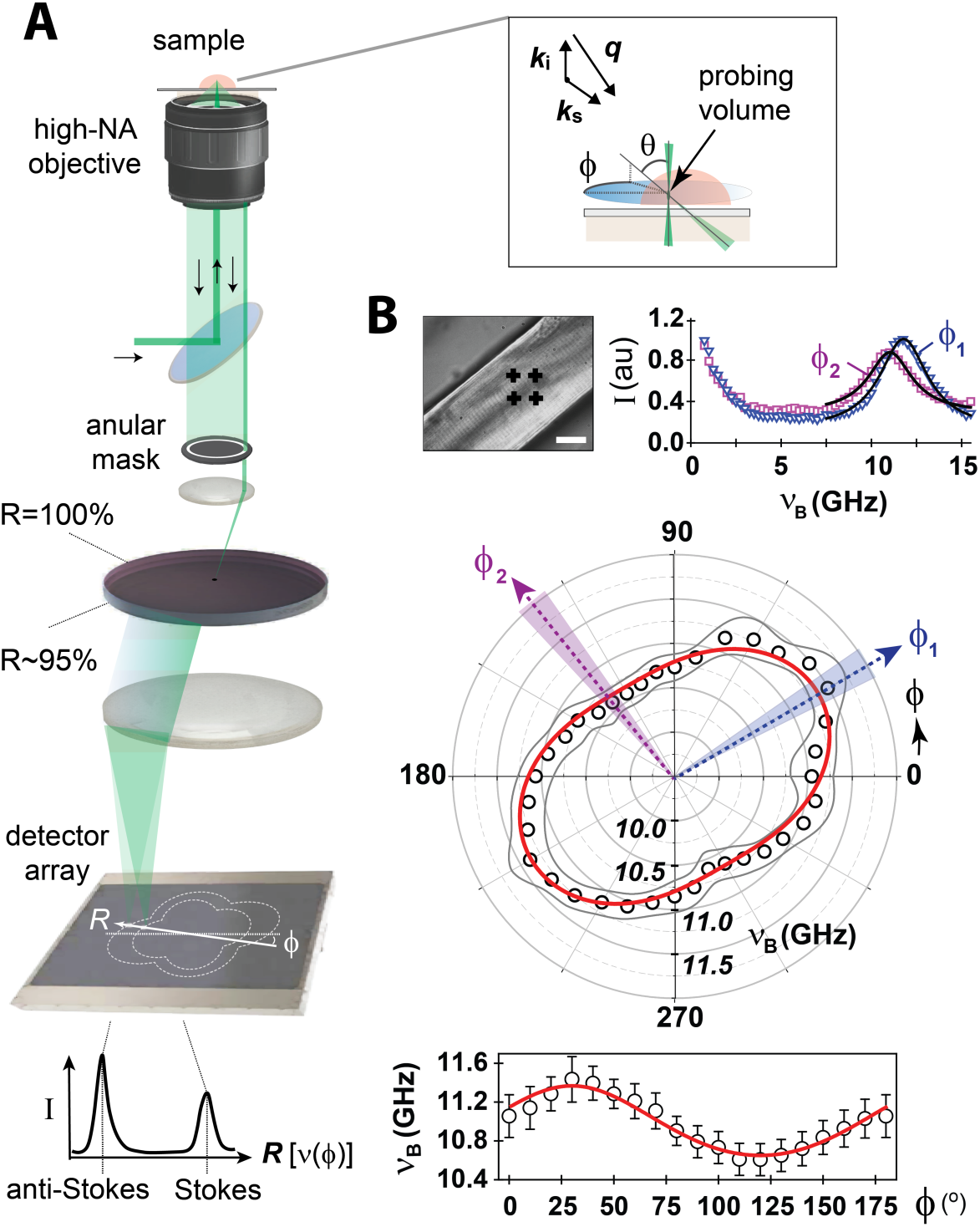
Brillouin Light Scattering Anisotropy Microscopy (BLAM). **(A)** Schematic of microscopy setup. *Inset:* probed phonon wavevector ***q*** for a given incident and scattering wavevector (***k*i** and ***k*s**). **(B)** Angle-resolved BLS frequency-shift *ϖ*B of muscle myofiber (red line=fit to transverse isotropic model). *Inset:* BLS-spectra in two azimuthal directions. Scale bar=25μm.

### Muscle fibers

We firstly tested Brillouin Light scattering Anisotropy Microscopy (BLAM) on isolated muscle fibers. The dependence of *ν*_*B*_ on the azimuthal angle *φ* (Fig 1B), are as expected from a transverse isotropic symmetry^17^, with the elastic moduli parallel and perpendicular to the fiber axis in good agreement with conventional confocal BLS-microscopy measurements using tilted excitation-detection angles on the same fibers (Methods). To test if BLAM can be used to monitor dynamic changes of cellular viscoelastic properties, we partially dehydrated the samples, observing an increasing anisotropy and average elastic modulus that reaches saturation quicker in the center of the fibers (Supplementary Fig 3), consistent with the expected outward water migration upon dehydration^18^. Changes in such dynamics may serve useful for assessment of muscle integrity from patient-derived muscle biopsies or muscle organoids and identification of compounds to treat muscular dystrophies or in aging-related muscle decline.

### Plant cell walls

Plant cell growth is understood to be primarily driven by turgor pressure, with asymmetric growth resulting from differential viscoelastic properties of cell walls^19^. The mechanical anisotropy of cell walls is thus crucial for determining growth and cell shape. It is determined by several factors, including the alignment of cellulose microfibrils^19^, as well as cross-linking of microfibrils and pectin^20^. The latter contribute to compressibility under fast deformations^21^ and are understood to lead to selective *softening* of walls^22^. Studies have suggested that turgor pressure may not always be necessary for establishing cell morphology in anticlinal leaf cells,^23^ and that a structural/phase transition of aligned pectin homogalacturonan (HG) nanofilaments upon demethylation causes a transverse expansion of filaments and *buckling* of cell walls. However the precise role that turgor pressure and the viscoelasticity of cell walls play in relation to cell growth and morphology from a material science perspective remains a much debated topic.^24^ Given the sensitivity to cross-linking and material phase state, BLS can offer unique insight into cell wall material properties underlying growth. While several studies have used BLS to study plant cell walls,^25–27^ these were not sensitive to symmetry-breaking in mechanical properties, which would be relevant for understanding growth mechanics, and can be probed with BLAM.

In Fig 2A we show representative measurements of *ν*_*B*_ for periclinal and anticlinal epidermal cell walls in *Arabidopsis thaliana* elongated hypocotyl cells as a function of azimuthal angle *φ*. The anisotropy is consistent with the mean orientation of constituent microfibrils and described by a transverse isotropic symmetry. The ratio of the maximum to minimum *ν*_*B*_ gives a measure of the degree of anisotropy (Fig 2B), and is significantly less than what is predicted for non-interacting aligned microfibrils (Supplementary Fig 4). This is consistent with the significance of cross-linking and the pectin matrix in the probed frequency regime, although can also result from a more complex microfibril network geometry^20^.

**Fig. 2.**
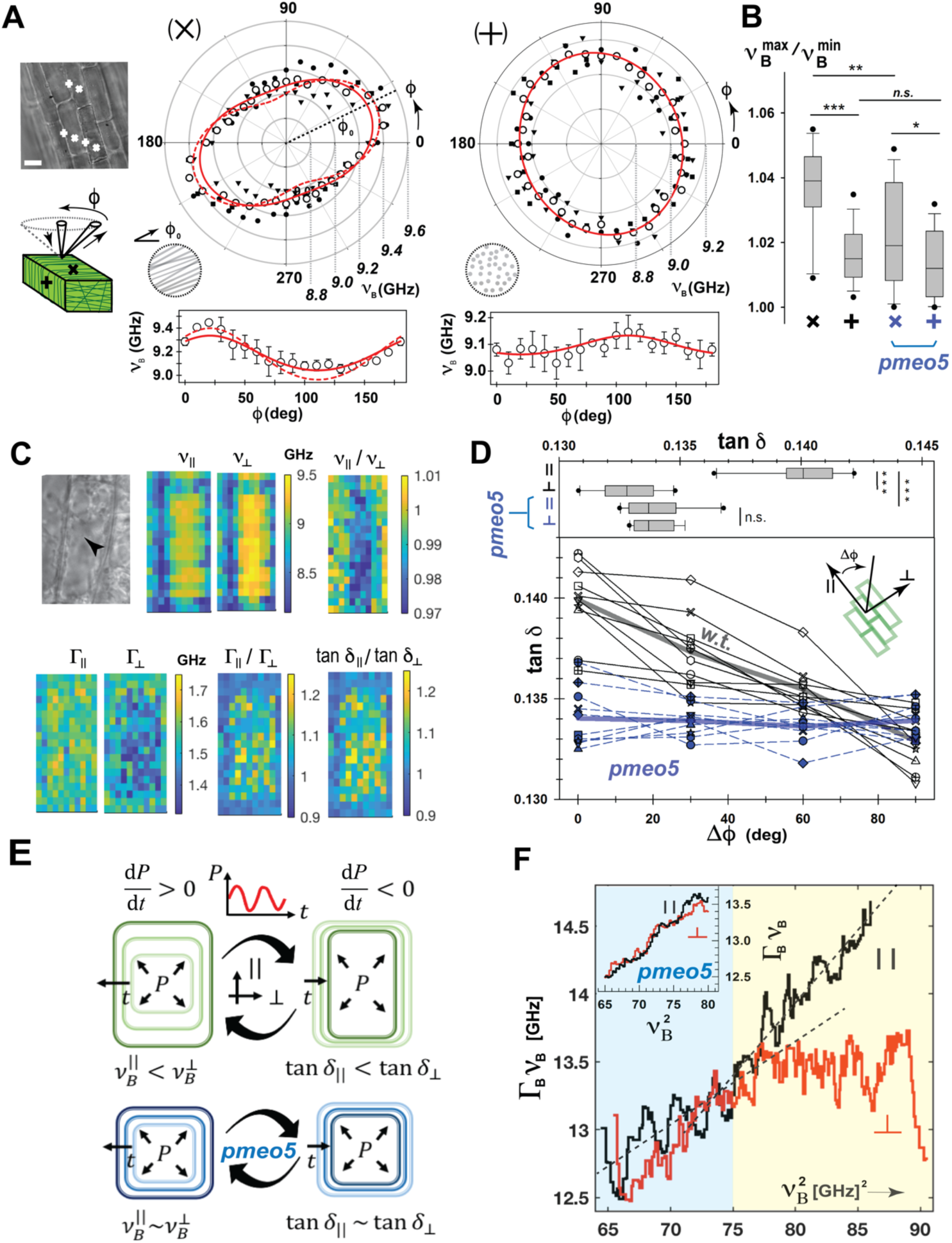
BLS anisotropy of *Arabidopsis thaliana* hypocotyl cell walls. **(A)** Representative plots of ϖB(ý) in cell walls perpendicular (×) and parallel (**+**) to growth axis. Solid line=fit to transverse isotropic model (Supplementary Text). **(B)** Ratio of maximum to minimum ϖB(ý) for cell walls with different orientations (n=10). **(C)** Representative maps of anisotropy in the BLS frequency-shift (ϖΒ), line-width (ρΒ), and loss-tangent (tan-8) for periclinal wall. **(D)** Comparison of anisotropy in tan-8 of periclinal cell walls for Wild Type (WT) and *pme5o* cells (n=6 each). (**E**) Schematic depicting anisotropic and isotropic growth as a result of fluctuations in turgor pressure, *P*. **(F)** Plot of loss (ϑ Γ_!_*ν*_*B*_) vs. storage (ϑ *ν*^2^_*B*_) properties of periclinal cell walls parallel and perpendicular to growth axis in WT cells. *Inset:* for *pme5o* cells. *p*\*\*<0.01, *p*\*\*\*<0.001, paired *t*-tests.

The relative contribution of elastic and viscous components can be gauged from the mechanical loss tangent, defined as: tan *δ* = Γ_!_/*ν*_*B*_, where Γ_!_ is the BLS-linewidth. tan *δ* can be understood as a measure of hysteresis upon microscopic deformations. It will by construction not be affected by anisotropy in the refractive index, which can be expected for cell walls, and which features in the relation between *ν*_*B*_ and elastic moduli (Supplementary Text). Spatial maps for periclinal and anticlinal walls (Fig 2C, Supplementary Fig 5), reveal a heterogeneous anisotropic landscape of tan *δ*, possibly associated with microscopic mechanical domains. For periclinal walls tan-8 is clearly larger (||, Fig 2D) indicating that the correspondingly altered *ν*_*B*_ is likely due to secondary structure on short time scales, rather than static structural anomalies in the microfibril alignment, and implies that local transient deformations are more *plastic* parallel to the cell-elongation axis.

We can investigate the effects of pectin demethylesterification via the inducible overexpression of pectin methylesterase (*pme5oe*)^23^. We find this results in a slight decrease in the average value of *ν*_*B*_, accompanied by a significant decrease in anisotropy (Fig 2B) and a reduction of the average value of the loss tangent in periclinal walls (Fig 2D).

How these observations might potentially affect plant cell growth can be appreciated in that growth occurs in periodic spurts,^28^ which can be expected to be accompanied by turgor pressure fluctuations due to *e.g.* calcium oscillations^28^ or environmental factors^29^. Our observations suggest that, during a phase of increasing turgor pressure, cell walls would incrementally be stretched more along the growth axis (since *ν*^||^ < *ν*^#^) resulting in an elongated cell shape. In a subsequent phase of decreasing turgor pressure, cell walls would however relax more completely perpendicular to the growth axis (since tan *δ*_||_ > tan *δ*_#_), and maintain an elongated geometry. HG demethylation would inhibit anisotropic and net growth, by virtue of walls expanding more isotopically and more effectively returning to their original dimensions following a turgor pressure drop. A schematic of this is shown in Fig 2E.

A plot of the viscous *vs.* elastic properties parallel and perpendicular to the elongation axis in periclinal walls (Fig 2F) show the anisotropy is due to a saturation in the viscous properties perpendicular to the elongation axis when the elastic properties exceed a certain value (indicated by blue-yellow shaded areas in Fig 2F). The decreased anisotropy upon pectin demethylesterification can be seen to be a direct consequence of a decreased elasticity (inset Fig 2F). That elevated values of tan-8 occur at isolated positions (Fig 2C, Supplementary Fig 5A), is consistent with cell wall loosening occurring at discrete locations^19^, and may also explain punctuated regions of increased BLS elastic moduli previously observed^26^.

### Mammalian cell nuclei

The eukaryotic cell nucleus is a highly dynamic heterogeneous and hierarchical structure. It resembles a complex polymer solution, consisting primarily of DNA folded into chromatin, together with a cocktail of other proteins.^30–34^ Chromatin is organized into spatial territories and domains with varying degrees of compaction, which serve to control gene expression^30^ as well as potentially other processes^33, 34^. It can exhibit both liquid-and solid-like behaviour,^31, 32, 35–37^ with dynamic phase separation known to play a crucial role^10, 31, 32^. While the BLS-measured elastic modulus has been shown to be sensitive to chromatin compaction^38^, to what extent this is locally anisotropic and how it varies spatially and dynamically can give valuable insight into the material state(s) of the nucleus, and static/transient structure.

Our measurements on live HeLa cell nuclei reveal a spatially heterogeneous elastic anisotropy, that span two symmetry motifs (Fig 3A) reminiscent of a transverse and orthotropic isotropic material. To quantify and present spatial anisotropy maps, we define the Brillouin Fractional Anisotropy (BFA) analogously to that employed in diffusion tensor analysis (which equals zero in the isotropic case and has a limiting value of unity), together with the vector *ȗ*. which points in the direction of the mean anisotropy (Methods). Figure 3C shows an example of a heatmap of the anisotropy and mean directions thereof in a live interphase HeLa cell. Compared to the nucleus, in the cytoplasm we find a significantly reduced anisotropy (Fig 3B) with less defined symmetry motifs. We attribute this to the averaging out of structure maintaining elements in non-polarized cells, compounded by the lower angle-averaged BLS frequency shift in the cytoplasm^39^.

**Fig. 3.**
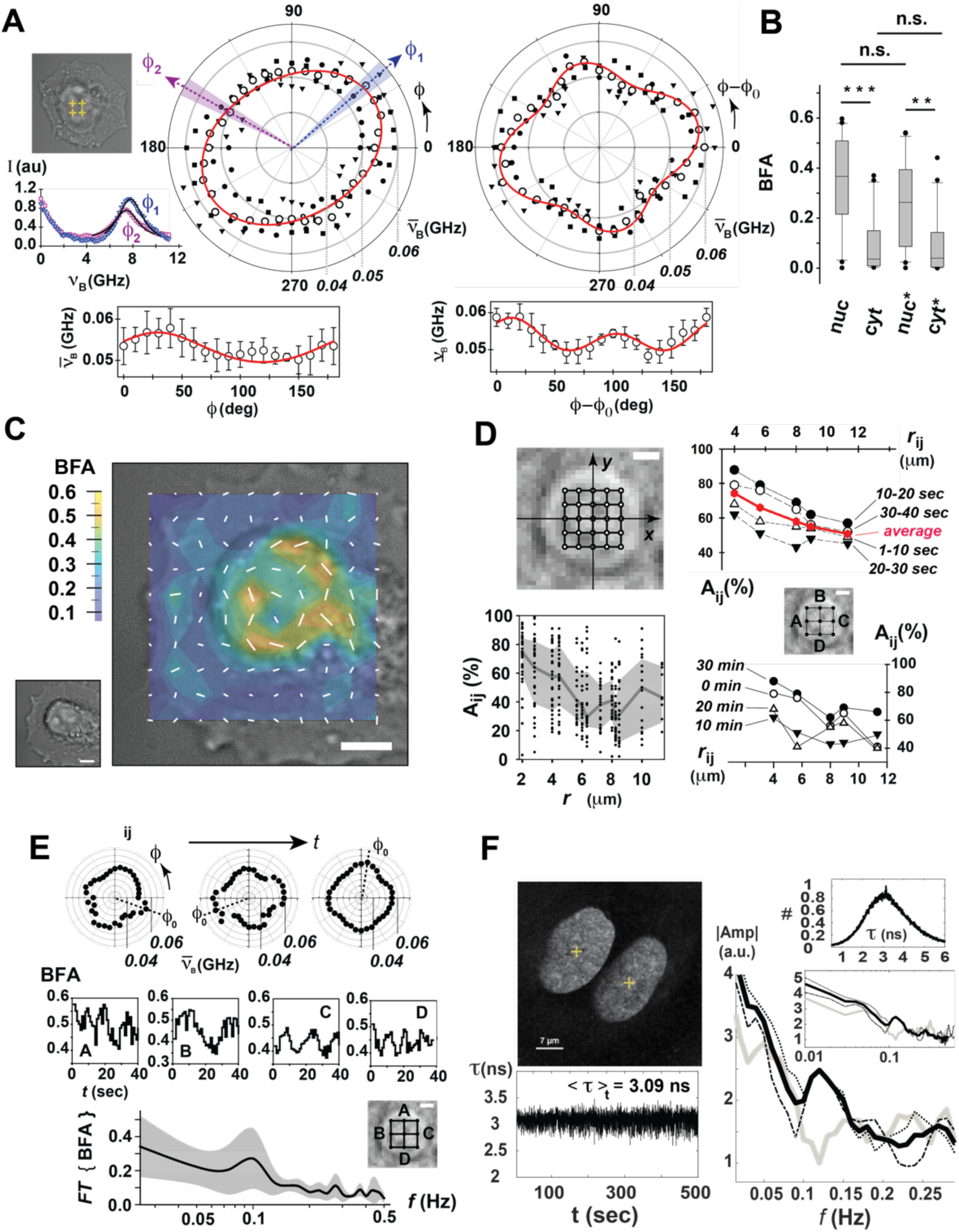
BLS anisotropy in cell nuclei. **(A)** Plots of two anisotropy motifs observed for the normalized Brillouin Shift *ν*_*B*_ in interphase HeLa cell nuclei (solid lines=transverse isotropic and orthotropic fit). *Inset:* representative BLS-spectra. **(B)** Brillouin Fractional Anisotropy (BFA) of HeLa cell nuclei and cytoplasm in live cells and cells during apoptosis (*). **(C)** Representative map of the BFA in a HeLa cell. Length and direction of the lines correspond to the magnitude of the BFA and mean orientation. Scale bar=4μm. **(D)** Alignment parameter *Aij* calculated from a scanned grid for interphase HeLa cell nuclei at different times. **(E)** BFA fluctuations at 4 positions and corresponding power spectrum (n=15 cells). **(F)** Time-trace of SIR-DNA fluorescence lifetime (ρ) in U2OS cell nuclei and associated power spectrum. *p*\*\*<0.01, *p*\*\*\*<0.001, paired *t*-tests.

In nuclei, we observe a finite nuclear-wide average anisotropy for both interphase HeLa and U2OS cells, with an inverse relation between the angle-averaged *ν*_*B*_ and the anisotropy, which increases in the early stages of apoptosis (Supplementary Fig 6). Given that this stage of apoptosis is associated with chromatin condensation^40^, and *ν*_*B*_ is larger for condensed chromatin^38^, this suggests condensed chromatin is associated with a lower anisotropy. Correlative fluorescence lifetime measurements of SiR-DNA stained nuclei reveal a decrease in the anisotropy is indeed correlated with a decrease in the SiR-DNA fluorescence lifetime (an indicator of chromatin compaction), Supplementary Fig 7. Consistent with anisotropy being associated with chromatin condensation, treatment with pro-apoptotic topoisomerase I inhibitor camptothecin, PARP1/2 inhibitor olaparib, cisplatin and hydrogen peroxide all notably reduced the BLS anisotropy, whereas chromatin decondensation induced by KDAC deacetylase inhibitor trichostatin A (TSA) result in an increased anisotropy (Supplementary Fig 6).

We also observe a positive correlation between the anisotropy and nuclear size (Supplementary Fig 6), likely due to smaller nuclei being in early stages of interphase with less decondensed chromatin. Interestingly, more asymmetrically shaped nuclei display a closer alignment between the mean anisotropy direction (*ȗ*.) and the nuclear elongation axis, as well as an overall larger nuclear-wide average anisotropy, suggestive of a connection between the BLS measured anisotropy and nuclear morphology (Supplementary Fig 7).

To quantify spatial anisotropy correlations, we define the parameter *A*_$%_as a measure of the similarity in *ν*_*B*_(*φ*) at two positions *x*_$_ and *x*_%_ (Methods). We find *A*_$%_ drops off with a characteristic distance of ∼2-5μm in interphase nuclei (Fig 3D), suggesting the existence of common structural motifs at such scales. Consecutive scans over the same areas show a change in the magnitude of *A*_$%_, but not so much the functional dependence, over the course of minutes (Fig 3D). The origins of these are revealed in time-series measurements at fixed positions which show periodic fluctuations in the anisotropy (BFA) with a characteristic frequency of ∼0.1 Hz (Fig 3E). The power spectrum of SIR-DNA fluorescence lifetime time-series measurements in interphase U2OS cells, show a similar enhancement at ∼0.1Hz that is suppressed in apoptotic cells (Fig 3F). This suggests that these may result from oscillations in the chromatin state.

The observed fluctuations are reminiscent of the ∼10 second lifetimes reported for transient chromatin cross-linking^35^, which were shown to cause local sol-gel transitions in interphase nuclei^36^. The two observed symmetry motifs (Fig 3A) may thus be indicative of the degree of cross-linking, which is consistent with the interphase nucleus existing in the vicinity of such a transition. Displacement Correlation Spectroscopy studies have also observed spatio-temporally coherent chromatin dynamics, driven by active nucleoplasmic streaming, with comparable spatio-temporal characteristics^37^. This suggests the observed dynamics are indeed associated with those of chromatin, which is not unexpected given the persistence length of chromatin, while ∼30nm at low frequencies^41^, will be larger at the probed high-frequencies^42^ and exceed the characteristic probed phonon wavelength.

Reliably obtaining spatial correlations of the anisotropy in nuclei requires fast grid scans, which are possible if one limits oneself to extracting the mean angular directions *ȗ*. (as opposed to the detailed anisotropic-motifs) by performing angular-binning (Methods). Figure 4A shows representative grid-maps of *ȗ*. for interphase U2OS cell nuclei obtained from grid scans performed in 9-10 seconds before and during apoptosis. These hint at potential spatial correlations, although the scan-times can be expected to average out any longer range spatial order in light of the above described fluctuations.

**Fig. 4.**
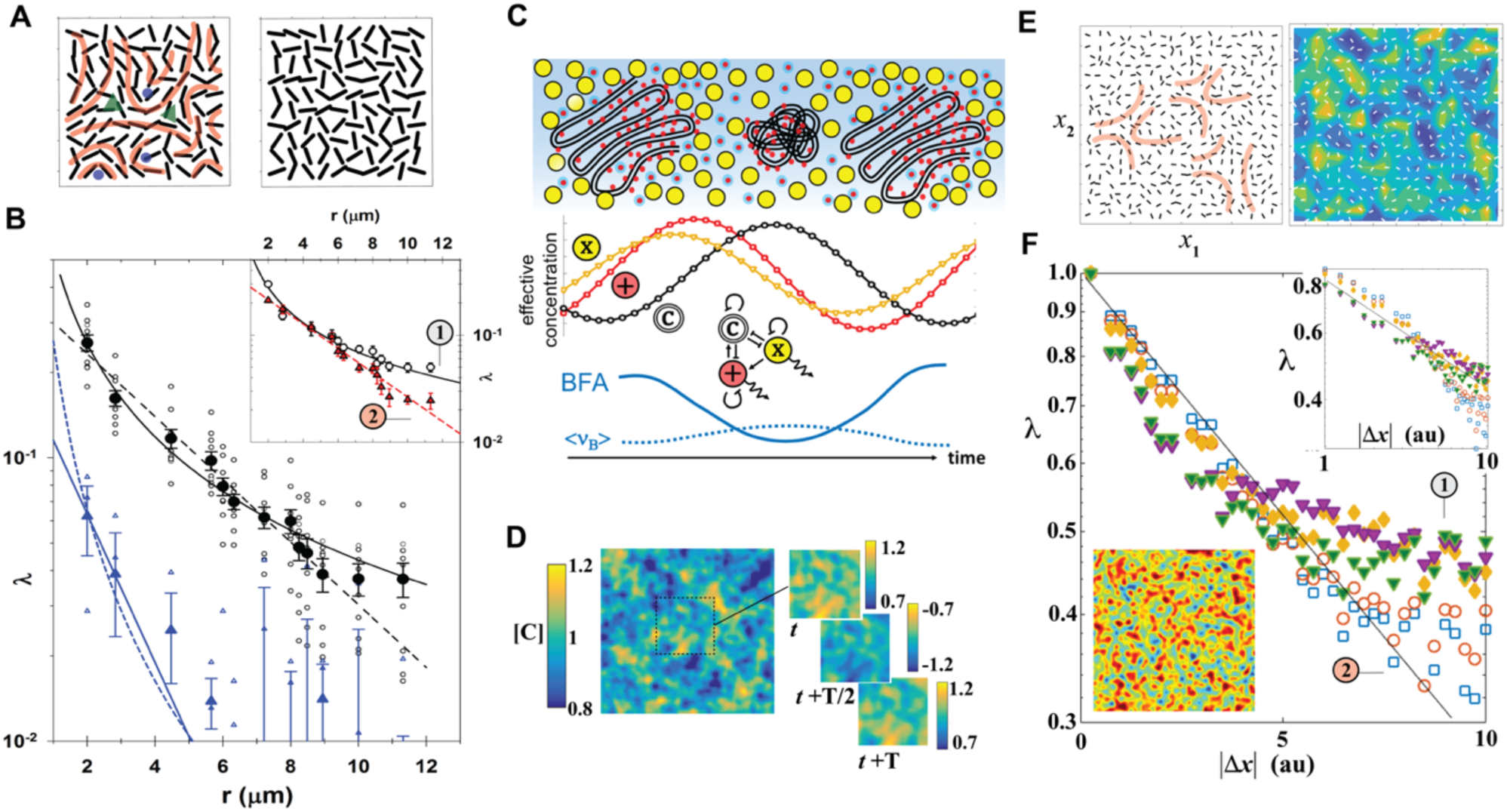
Spatial and temporal correlations of BLS anisotropy in cell nuclei. **(A)** Representative map of *ȗ* in interphase HeLa cell nuclei before (left) and during (right) apoptosis (step size=1μm). **(B)** Orientational correlation function for nuclei [O=live interphase (•=mean), **l::,** =following apoptosis (**•**=mean)]. *Inset:* Same data sorted into two subsets with high (O) and low (**•**) short distance correlations, with power-law (dashed line) and exponential (solid line) fits respectively. **(C)** Schematic of 3-species model (red=cations, yellow=crowding agent, black/white=condensed chromatin), showing calculated temporal evolution of species concentrations and BFA for stable Spatio-Temporal Oscillatory Solution (STOS). **(D)** Example spatial map of condensed chromatin concentration [C] for a STOS with period T. **(E)**. Quiver plot of electric potential gradients superimposed on condensed chromatin concentration for a STOS. **(F)** Normalized orientational correlation functions (/\) for two different reaction parameters (solid and open symbols) showing power-law and exponential -like scaling (*Top inset:* on log-log scale). *Lower inset:* snapshot of condensed chromatin concentration from simulation.

We overcome this by performing coarser (2μm-grid) nuclear-wide scans in < 3 seconds, from which we can calculate an orientational correlation function λ (Methods) and identify potential spatial correlations (Fig 4B). Here an exponential scaling of λ would suggest largely uncorrelated dynamics, whereas an algebraic (power-law) scaling can be considered indicative of the existence near a critical state^43^. We find λ can be described well by both an exponential and an algebraic function (Fig 4B). However, when sorting cells into groups with larger and smaller λ at short distances (Supplementary Fig 8), we find the former and latter are statistically better described by an algebraic and exponential scaling respectively (inset Fig 4B). This suggests that a fraction of cells are at a critical state (ordered-disordered transition).

To gain insight into the molecular origin of these observations we note that a key parameter for describing chromatin dynamics is the solvent-chromatin *friction*^37^, which is dictated by the effectiveness with which cations dynamically screen chromatin and decouple chromatin-solvent motion, and which the high-frequency regime is acutely sensitive to^6^. In addition to decoupling chromatin-solvent dynamics, cations will also screen chromatin self-interaction, typically promoting chromatin condensation^44^. Evidence that we are looking at more coupled *vs.* decoupled chromatin-solvent dynamics for more condensed *vs*. decondensed chromatin states can be seen by looking at the scaling of the high-frequency elastic modulus with respect to condensed chromatin fraction (Supplementary Fig 9, Supplementary Text). Various larger molecular species in the nucleus are known to also promote chromatin condensation and decondensation (Supplementary Text), often through active processes, but also as a consequence of molecular crowding^45^.

Based on the above, we can construct a 3-species reaction-diffusion model (Fig 4C) where condensed chromatin fraction “C” is promoted by mobile cations “+”, and inhibited by a larger diffusive species “X” (which may serve as a crowding agent), to try explain our observations (Methods). Such a model can support Spatio-Temporally Oscillating Solutions (STOS) manifested as oscillating Turing patterns of the different species (Fig 4D, Supplementary Text). To calculate anisotropy maps representative of *ȗ*. we note chromatin can be expected to orientate itself according to the local electrostatic potential gradient (owing to the negatively charged DNA backbone) which can be calculated from spatial variations in the cation concentration (Fig 4E).

Calculating *λ* from these simulations (Methods) we find that, depending on the choice of activation/inhibition rates, one can obtain algebraic-or exponential-like scaling (Fig 4F), mirroring what is observed in cell nuclei (Fig 4B). From the parameter constraints required to realize stable STOS we can estimate the effective size of X to be on the order of ∼50kDa, which fits with several ATPases and Hystone Acetylasetrasferases known to cause chromatin relaxation/decondensation (Supplementary Text). As with most reaction-diffusion models, activation/inhibition rates need to be tightly regulated for stability, although solutions are found to be stable to small amounts of noise if one includes a weak restoring force for each of the rate-constants (Supplementary Fig 10), which can presumably be provided by active processes.

## Discussion

We have introduced a setup for imaging the high-frequency viscoelastic anisotropy in living soft-matter (BLAM), and presented a cross-section of what it can reveal in different cellular components. In plant cell walls our results show that BLAM can observe both the elastic and viscous anisotropy of cell walls. A model of how this may affect asymmetric cell growth via a “ratchet” mechanism is presented, where turgor pressure fluctuations are responsible for *establishing*, but not *maintaining*, elongated cell shape, and asymmetric growth can be modulated by pectin demethylesterification. The relation between the elastic and viscous properties (Fig 2F) can be explained by having distinct viscous relaxation times parallel and perpendicular to the growth axis (Supplementary Fig 6), with the amount of methylation dictating the degree of anisotropy.

In mammalian cell nuclei we have found a surprisingly rich dynamic viscoelastic landscape, that hints at the nucleus existing in the vicinity of an ordered-disordered state with long range correlations. We propose a simple model that explains our observations as a combination of faster and slower diffusive processes creating a dynamic electrostatic landscape that modulates the condensation state of chromatin. While a significant portion of the model relies on *passive* phenomena, active processes would be required for maintaining STOS in the presence of dissipation, as well as being potentially also involved in the inhibitory action (chromatin decondensation) of the slower diffusive species. While there are also passive mechanisms that can serve for the regulation of rate constants (such as pH-sensitive membrane porosity^46^), it is likely these too are aided by active processes, the disruption of which result in changes in the electronegativity and break down of the STOS state. The observed suppression of oscillations in apoptotic, cisplatin-, and etoposide-treated cells, which are all associated with increased acidity and chromatin condensation, can be understood as the result of a disruption in this regulatory mechanism.

Our results on the nucleus also address its both liquid-like and solid-like properties. The scaling of the orientational correlation function, together with the oscillatory anisotropy, are suggestive of the nucleus existing near a critical order-disorder transition, with behaviour characteristic of a quasi-nematic liquid crystal. There is a functional appeal of existing in such a state, characterized by long-range spatial correlations, that can be reconfigured rapidly with minimal energy expenditure for dynamically coordinating distant nuclear processes, however the precise biological significance is yet to be explored.

With minor modifications, the employed setup should also be able to study the anisotropy of the shear modulus (via the scattering from transverse acoustic phonons), however we have not been able to do so in our current implementation, owing to the smaller energies and scattering cross sections of these modes. To this end, as well as for extracting faster temporal correlations, approaches employing stimulated BLS^47^ may prove useful if the geometrical constraints can be overcome.

In conclusion, we have shown how BLAM can be used to measure the structural anisotropy in biological matter. We have also shown that it can give us a unique glimpse into the dynamic interplay between the microscopic and the macroscopic worlds in active soft matter and thereby the origins of structure and emergent phenomena associated with the probed mesoscopic scales^48^. Applications of BLAM can be expected to span material engineering and characterization, medical diagnostics and quality control. Studies of transient broken rotational symmetry also have the potential to uncover new universality classes for phase transitions^49^ and identify novel active material states^50^.

## Acknowledgments

We acknowledge support from the Vienna Biocenter Core Facilities (Plant Science Facility and Advanced Microscopy Facility), and thank Alex Dammermann for critical reading of the manuscript. We acknowledge funding from the EU Interreg grant V-A AT-CZ, RIAT-CZ, ATCZ40 (HK, DC, MU, KE), a Marie Skłodowska-Curie Action postdoctoral fellowship (HK), the City of Vienna and the Austrian Ministry of Science -Vision 2020 (KE), and the Austrian Science Fund -FWF P34783 (KE).

## Author contributions

Conceptualization & project administration: KE. Methodology: HK, DC, MS, LS, LMA, MU, IY, DS, WJW, AP, JP, KE. Investigation & visualization: HK, DC, MS, LS, LMA, KE. Funding acquisition: HK, MU, KE. Supervision: MU, IY, DS, JP, KE. Writing – original draft: KE, IY. Writing – review & editing: All Authors

## Data and materials availability

Data for this study are available in the main text or supplementary materials, and where not the case, will be made available publicly upon publication.

## Supplementary Materials

Supplementary Fig.’s 1-10, and Supplementary Text

## Methods

### Microscopy and Spectroscopy

#### Angle-resolved Brillouin Light Scattering Microscopy

Angle-resolved Brillouin Light Scattering measurements were performed on a custom built microspectroscopy setup. A schematic detailing the main features of the setup used for live cell studies is presented in Fig 1A and in more detail in Supplementary Fig 1D (setup “A”). An alternative version of the setup used for measurements on muscle fibers and plant cells is shown in Supplementary Fig 1E (setup “B”). A key component is the modified Virtual Imaged Phased Array (VIPA) which serves as the dispersive element. In our design, the Cartesian degree of freedom of a conventional VIPA is substituted with an azimuthal degree of freedom, by replacing the VIPA entrance-slit with an entrance hole (Supplementary Fig 1A-1C). We refer to this component as the radial-VIPA owing to the radial spectral projection rendered. Details on its fabrication and alignment can be found in the Supplementary Text.

In both cases (setup “A” and “B”) a single frequency laser (750mW, 532nm, Torus, Laser Quantum) with an optical isolator was used. In an earlier setup used for muscle myofiber measurements and several of the plant measurements a 561nm (Cobolt Jive) laser was used which was later replaced for technical reasons. This yielded identical results for the plant cell walls when correcting for the different wavelength, and slightly different Free Spectral range of the radial-VIPA. Despite all components being either broad band anti-reflection coated or having a significant transmission at both wavelengths, the BLS signal was notably weaker for 561nm likely due to the smaller scattering cross-section.

A small portion of the light is coupled out to a photodetector by a glass plate to indirectly assess the laser power at the sample (via a look-up table) and monitor laser stability, as well as to be passed through a gas absorption cell to assure an absorption line is tuned to the laser (see below). For setup “B” the light is then passed through a pair of axicons (physical angles 20° and 40°) as well as an S-plate to create an annular profile with radial polarization^51^. The laser light in both setups is subsequently coupled into the main optical path via a Non-Polarizing 90:10 (T:R) Beam Splitter (NPBS).

For setup “B” the detected light is incident on a custom machined blacked thin metal disk with a single hole (diameter 800μm) a distance of 4.5mm from the disk center and rotation axis (Supplementary Fig. 1E) that is mounted on a manual *xy*-stage (*z* being the optical axis). The disk is thereby aligned such that its center coincides perfectly with the subsequent objective lens. The position of the axicons are adjusted to create an annular profile that coincides with this hole at all rotation angles. During imaging the disk is rotated via a rubber belt connected to an electric dynamo motor (extracted from a toy RC car) at >2000 rpm (rotations per minute).

The light in both setups is subsequently focused onto the sample with a 1.49NA/100x immersion-oil objective lens (UAPON, Olympus) mounted in an inverted geometry. For setup “A” this will correspond to normal incidence, whereas for setup “B” this will correspond to an incidence of approximately 77 degrees relative to the optical axis (*z*). The effective Numerical Aperture (NA) of the probing beam is in both cases approximately 0.2 and results in a measured focal spot with Full Width at Half Maximum (FWHM) of 1.3-1.4 microns.

For setup “A”, the back-scattered light from the sample, after passing through the NPBS passes through an annular (ring) mask. This was selected from a plate of >20 masks (406 aluminum coated quartz, custom made by Photo-Sciences Inc., USA) that had inner diameters ranging from 3-5mm and annular thicknesses from 0.5-1mm: the effective detection NA thus being between 0.1-0.2. These would allow for the mean detection angles between 52 and 90 degrees relative to the probing light for a sample of refractive index 1.4. The particular annular mask was chosen to be slightly below that which would result in total internal reflection (which was discernable from visual inspection) to maximize the polar angle and thus in-plane anisotropy measurable. For each sample the mean angle was calculated based on the slit width, which was used in any quantitative analysis for calculating *e.g.* stiffness tensor components (see Supplementary Text). A flip-mirror was used to optionally couple all the detected light to a 1GHz scanning Fabry Perot etalon (Toptica, FPI 100-0500-3V0) to measure the elastic scattering peak and assess laser stability.

The elastic scattering peak from samples, used for spectral deconvolution (see below), was measured by detuning the laser from the absorption cell resonance slightly, inserting neutral density filters in the beam path and measuring under all the same acquisition settings. This was only done once at the beginning and end of each sample measurement when relevant at a position manually chosen to be roughly in the middle of the measured area.

The light was in both setups subsequently passed through a heated Iodine gas absorption cell adjusted to have an absorption line coinciding with the elastic scattered light. This was achieved by outcoupling a small portion of the laser light, passing this through the absorption cell, and weakly focusing this on a Si-photodetector (Supplementary Fig 1D). The best possible elastic suppression was achieved by finely tuning the temperature of the gas cell, and dynamically adjusting the laser current (which results in a small spectral shift of the laser emission frequency) in a simple feed-back loop.

The detected light is subsequently focused through a 150 μm pinhole (corresponding to ≈0.7 Airy Units (*AU*) for our setup). The light is then collimated and focused with a *f*=250 mm aspherical tube lens through a hole in an (45°) off-axis 2” parabolic *f* = 200 mm mirror mounted on a six-axis fine tip/tilt stage, and into the 100μm entrance hole of the custom fabricated radial-VIPA (see Supplementary Text) also mounted on a similar six-axis high-precision alignment-stage designed for fiber coupling (Thorlabs, DE). For the designs employed in these experiments, etalon thickness of 3mm were typically used (corresponding to a free spectral range of ∼33-34 GHz at the probed wavelengths). We employ a reflective VIPA configuration (*i.e.* the entrance hole is on the R=95% reflective coated side), which significantly reduces the physical footprint of the setup (that is < 1 meter in length). Doing so does not affect the dispersion characteristics of the VIPA, which can be described using the same paraxial-approximation relations as that of a single conventional VIPA^52^. Details pertaining to fabrication of the VIPA are described in the Supplementary Text. The choice of collimating lens and focusing lens (Supplementary Fig 1D and 1E) were chosen such that the effective point spread function at the entrance of the radial-VIPA was ∼90μm (*i.e.* slightly smaller than the diameter of the entrance hole).

The dispersed light from the VIPA is focused by the parabolic mirror onto an intermediate image plane, prior to which it is passed through a custom fabricated radial apodization filter (Supplementary Text). An iris is placed in this intermediate image plane to mask out high orders. The light is re-imaged (with a magnification factor of 2) onto a cooled EM CCD Camera (Andor, iXon), with a second iris placed in the intermediate Fourier plane effectively serving as a Lyott stop to partially filter out higher frequency interference signals from the sample and precluding optics^53^. The entire setup was controlled by custom Labview (National Instruments) and Matlab (Mathworks) based scripts, which also included moving the piezo-stage on which the sample was mounted laterally in a grid and acquiring the spectral projections at each position. For the case of the fast grid-scans (Fig 3) the magnification on the EM-CCD camera was reduced (by exchanging the final tube lens and camera position) such that the peak width was reduced to occupying ∼2-4 pixels on the camera.

Samples were all prepared on high-precision glass bottom dishes (Matek) and measured from below (*i.e.* through the cover glass). The glass bottom dishes are mounted on a custom holder fixed on top of a long travel-range piezo stage (733.2CL, Physik Instrumente, DE) which is in turn mounted on a manual translation stage (Olympus, JP) that allows for course positioning.

#### Spectrometer alignment & calibration

The alignment of the spectrometer was such that the energy was concentrated primarily into a single order, and the anisotropy was fitted to the most prominent Stokes peak in the BLS spectra. The reason for this is to maximize acquisition time/throughput (by maximizing the photon count for a single BLS peak) as well as avoiding potential distortions caused by the non-perfect Iodine gas cell absorption spectra (which was found to distort the anti-Stokes peak shape). The former proved important for measuring sensitive samples (such as live cells), however, also requires more rigorous calibration measures. The spectra of a known isotropic liquid or else the elastic scattering light (which should both trace an isotropic projection) were used to correct for subtle aberrations in the BLS spectra on the fly using a polynomial correction factor for each azimuthal angle increment, which was verified to be adequate and applied to all measured BLS spectra (Supplementary Fig 2). Prior to each experiment the tip-tilt and position of the radial-VIPA were finely adjusted to qualitatively minimize such aberrations. Calibration spectra obtained from isotropic materials (water and immersion oil) on the edge of the same cover slip and below the sample, were used before and between measurements to register the spectral dispersion axis, using the dispersion relation for a single-axis VIPA in the paraxial approximation^52^ and a two parameter fit (in the same way as for a conventional VIPA). The lateral spatial resolution of the setup was measured to be ∼1.5μm by scanning across a plastic/methanol interface (Supplementary Fig 2). This is consistent with the expected optical resolution when one considers the convolution of the excitation and detection optical transfer functions, and the relevant acoustic scales in both setups. The spatial resolution for our setup can be assumed to be limited by the optical resolution which exceeds the characteristic spatial scales of the probed phonons. All grid scan steps were larger than this resolution to avoid spatial correlations due to Point Spread Function (PSF) overlap between positions. For setup A, the axial resolution is comparable to the lateral resolution, owing the crossing of the excitation and detection PSFs which yield a significantly smaller effective PSF (see inset in Supplementary Fig 1D).

#### Experimental conditions

All measurements were performed with maximum EM gain settings of the camera (iXon, Andor). For the myofibers and Arabidopsis cells the laser power at the sample was ∼7-10 mW at 532 nm (and <20 mW at 561 nm) and the acquisition times (per spectral projection) were <1 second. For the HeLa and U2OS cells the laser powers were ∼5-6 mW and at all times < 8 mW (at 532 nm) and the acquisition time <100ms. The higher powers and longer acquisition time for the Arabidopsis were necessary to reliably extract the peak width. In all cases it was confirmed that these laser powers did not adversely affect the samples by inspecting samples ∼5-10min after measurements with the widefield camera, and looking for any morphological anomalies, signs of blebbing, *etc.*. We note that the overall sample laser exposures are significantly less than routinely employed in conventional BLS microscopy (which have been shown to not cause any significant phototoxicity in these same samples), due to the sparse/coarse grid scans we employ. Furthermore, due to the effective NA being lower than that commonly used in high-NA confocal Brillouin imaging, the photon flux at a given position is also significantly less. All measurements were performed at room (lab) temperature, T=23(+/-< 2) C, with comparative measurements checked to be sure temperature variation was less than 0.5 C.

#### Fluorescence measurements

High resolution fluorescence confocal measurements (to obtain the morphology and average fluorescence intensity of U2OS cell nuclei) were performed on a Microtime 200 RAPID-FLIM setup (Picoquant, DE), with a 1.4 NA/60x water immersion objective-lens (Olympus). A 640 nm laser diode operating in Continuous Wave (CW) (for steady-state measurements) and 40 MHz pulsed (for fluorescence lifetime measurements) was coupled into the system and used for excitation with suitable quad-pass dichroic and bandpass filters for detection. For the fluorescence lifetime time-traces the laser was parked at a point in the nucleus and data was acquired for 500 seconds. The data was divided into 10 ms second bins, and the lifetime for each bin was obtained by fitting a single exponential after deconvolution with the measured instrument response function. *X*^&^ values were typically around 1+/-0.1, and gave lifetimes between 2.9-3.15 ns. All Fluorescence lifetimes were fitted with Symphotime (Picoquant, DE). Fourier transform analysis of the lifetime traces were performed using a custom written Matlab script. For correlative fluorescence BLS measurements part of the signal after the pinhole was coupled out using a short-pass dichroic (cut-off 535 nm) and focused onto a Photo-Multiplier

Tube (PMT) (Thorlabs, DE). Fluorescence data was acquired in series for each voxel using a custom Matlab acquisition script adapted from that developed for a previous correlative fluorescence-BLS microscope^26^. Though for the BLS measurement this excitation wavelength is quite far off from the absorption peak of SiR-DNA, the emission was confirmed to originate from SiR-DNA fluorescence by performing spectrally resolved measurements with a fiber-coupled QE-Pro Ocean Optics fluorescence spectrometer in place of the PMT.

#### Widefield images

Widefield reflected white light images were obtained on the BLS setup using a 40x long working-distance air objective (Olympus) on top of the instrument, and imaged on a sCMOS camera (FLIR) with a 250 mm tube lens (effectively increasing the magnification to ∼50x on the pixel array, and achieving a lateral resolution comparable to that of the regions probed in BLS anisotropy point measurements).

#### Conventional BLS measurements

High incident-angle conventional BLS measurements on myofibers were performed by redirecting the light after the pinhole with a flip mirror, coupling this into a single mode fiber, and running this to a conventional 2-VIPA BLS spectrometer (described previously^26^). To probe the predominantly parallel and predominantly perpendicular component of the mechanical properties, a makeshift blacked plastic mask (with 0.3-0.5 mm hole), effectively blocking all but a portion of ∼10 degrees of the excitation and scattered light at large excitation angles, was placed immediately behind the objective lens and manually rotated around the optical axis to obtain measurements at different azimuthal angles. An angular correction factor was calculated using an isotropic sample (distilled water) in the same manner as for the radial-VIPA (see above). Subsequently measurements at 90° degree increments predominantly “parallel” and “perpendicular” to the myofiber axis were made and corrected with the alignment factor. The fiber axis was identified from widefield transmitted white light images. The laser power was significantly increased but still less than ∼7 mW at the sample. Acquisition times were 1-2 seconds and averaged over 9 positions of a fine 0.2 μm spaced 2D grid. The effective values of the elastic modulus parallel and perpendicular to the fiber axis was calculated in the same manner as for the radial VIPA (as described in Supplementary Text) using Eqn. S13 and S14. The refractive index and density for the myofiber were taken to be *n*=1.38 and *ρ* =1.06 kg.m^-3^ respectively. The refractive index for these calculations was assumed to be isotropic (although this is atrictly speaking unlikely to be the case). This, or the fact that the true values of the refractive index deviated from those assumed, will not affect the comparison between the two spectroscopy approaches, which is the purpose in the current context. For a hydrated myofiber we obtained *c*_#_ = 8.31(±0.47) GPa and *c*_||_ = 12.89(±0.55) GPa using the single high-incident angle confocal BLS, compared to *c*_#_ = 8.17(±0.51) GPa and *c*_||_ = 13.40(±0.38) GPa for the same fiber using the radial-VIPA setup.

### Sample preparation

#### Myofibers

EDL muscle was surgically removed from adult (6 month old) mice. The muscle was than digested with collagenase (type1, 2mg/ml, Sigma) supplemented in media (DMEM with 10% FCS and penicillin, Life Technologies, final conc. 50 U/mL of penicillin) for 2 hours at 37° C. After digestion, individual fibers were released from the muscle by gentle trituration. Healthy elongated myofibers were separated from the debris and transferred to high-precision glass bottom dishes containing full media for measurements. For dehydration studies excess media was removed and the fibers were allowed to dry under ambient conditions (∼23C, ∼50% air humidity) while on the microscope stage.

#### Arabidopsis thaliana

The Arabidopsis thaliana seedlings ecotype Wassilewskija (WS) were grown on MS media^54^ supplemented with 1% sucrose and solidified with 0.8% agar at standard growth condition (22°C and 16 hour light / 8 hour dark cycle). Measurements were done in roots of 7-day old control seedlings (WS) and seedlings overexpressing the pectin methylesterase 5 (PME5oe) as described in^55^. The ethanol induction was performed by adding ≈100 ml of ethanol to the bottom of the MS plate that were left standing vertically for 18 hour in the plant growth incubator.

#### HeLa and U2OS cells

U2OS and HeLa cells were maintained in DMEM supplemented with 10% heat-inactivated FCS, 1 mM L-glutamine, 100 U/mL penicillin and 100 μg/mL streptomycin. For imaging, the cells were grown on uncoated high-precision glass-bottom MatTek dishes. U2OS cells were when relevant subsequently stained with 1 μM SiR-DNA (SiR-Hoechst; lambda abs 652/em 674 nm). Measurements in the early stages of apoptosis were taken as those immediately prior to the first observations of blebbing. U2OS cells were treated with 10 µM trichostatin A (TSA) for 6h (AbMole M1753), 25 µM camptothecin for 1h (Sigma-Aldrich C9911), 20 µM etoposide for 1 hour (Sigma-Aldrich E1383), 30 µM cisplatin for 2h (Sigma-Aldrich 479306), 10 µM olaparib for 2 hours (Selleck Chemicals AZD2281), 200 µM H2O2 for 10 min (Sigma-Aldrich 216763) and 1 mM temozolomide for 4h (MedChemExpress HY-17364). DMSO was used as a control at 0.3% (Sigma-Aldrich 472301).

## Data Analysis and Modeling

The angular spectral projections were first interpolated into a set of 359 linear projections (corresponding to radial profiles of the measured projections for each degree). These were binned into 10 degree splices each composed of the average of the 10 constituent spectra. For the very fast grid scans these were initially interpolated into 35 projections and then averaged into 90 degree splices (4 quadrants). The binned spectra were subsequently corrected for aberrations based on measurements of known isotropic samples (Supplementary Fig 2). Knowledge of the Free Spectral Range (FSR) and measurements of two reference spectra (ethanol and distilled water) in the same imaging session were used for spectral registration. The spectra were deconvolved (performed in Matlab by converting spectra into Fourier space^56^) with the shape of the elastic scattering peak, measured in the same sample–see above. Non-linear least square fitting of deconvolved spectra was performed using the Matlab function *PeakFit* (as described in *e.g.*^26^) with a Lorentzian function and a linear background, typically in the range between ∼4-15 GHz, to extract the BLS peak position, as well as the peak width for the case of the *Arabidopsis thaliania* samples. The elastic or loss moduli of interest were when applicable calculated based on the scattering angle as described in the Supplementary Text.

The *Brillouin Fractional Anisotropy* (BFA) was defined in an analogous manner to what has previously been used in angle resolved Optical Coherence Elastography^57^ as:

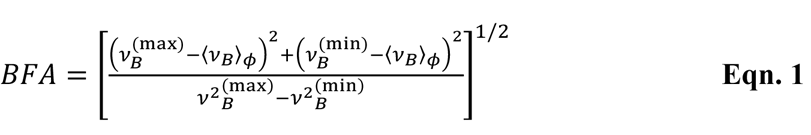

where *v*(^*max*)^ and *v*(min) are the maximum and minimum values of *ν*_*B*_(*φ*), and 〈*ν*_*B*_〉_8_ = *v*^)/^ ∑_9_ *ν*_*B*_(*φ*_9_) is its angular average (*N* = number of spectral projections). The *normalized*

*Brillouin frequency-shift*, *v*_!_(*φ*) is defined as in^12^:

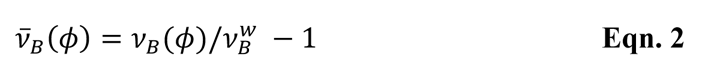

where *v*: is the frequency shift of distilled water measured with the same setup and in the same experimental session. *v*_!_ is a measure of the relative deviation of the spectra from that of water, and as such is a useful metric for presenting BLS data in samples with high water content, where the deviations therefrom are often subtle.

The mean direction and magnitude of the BLS anisotropy (as presented in *e.g.* the quiver plots of Fig 3 and Fig 4) are defined at each point by the vector:

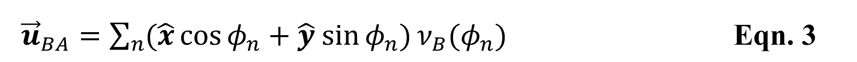

The angle *φ*_9_here represents the midpoint of the respective splices. This vector is also used to calculate the *alignment parameter A*(*r*) (plotted in Fig 3D), between two positions in a grid scan which was defined as:

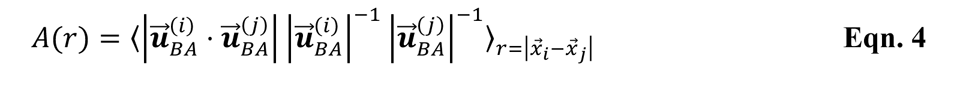

The orientational correlation function (plotted in Fig 4) was calculated using:

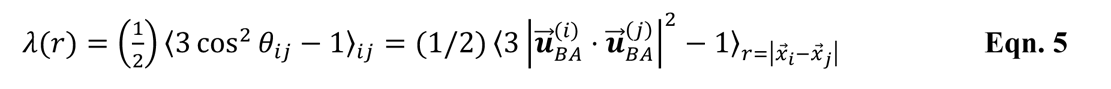

where *θ*_$%_ is the angle between the mean anisotropy direction at *x*_$_ and *x*_%_.

Calculations of Voigt and Reuss effective responses used the following equations respectively:

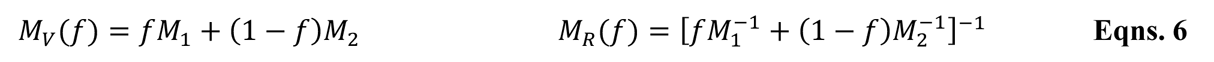

where *M*_/_and *M*_&_are the moduli of the two components and 0 < *f* < 1 is the filling factor. These also serve as the basis for modeling the transverse anisotropic properties of muscle fibers and cell walls, and described in the Supplementary Text.

The fit for the Reuss-Voigt model (Supplementary Fig 9) additionally assumed the following (see corresponding discussion in Supplementary Text):

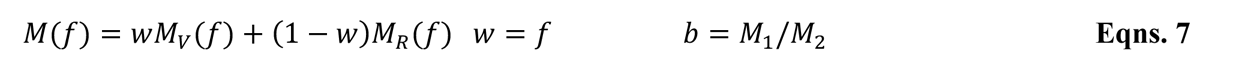

To simulate the processes illustrated in the reaction-diffusion model shown in Fig 4C, a 3 species activator/inhibitor model was assumed (*inset* of Fig 4C). Specifically, 3 differential equations were constructed of the form:

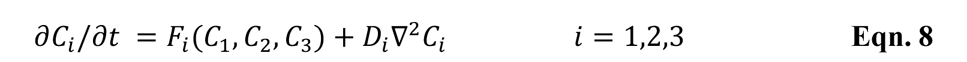

where *C*_/_, *C*_&_ and *C*_C_ represent the fraction of condensed chromatin, cation concentration (that promote chromatin condensation), and an additional species that inhibits chromatin condensation and also serves as a crowding agent, respectively. *D*_$_are their respective diffusion coefficients, and *F*_$_ describes the reaction/interaction terms (see Supplementary Text). The evolution over time of the concentrations are solved on a 200 x 200 2D discretized space in Matlab (Mathworks, DE), and ran for >500 time point (for the coefficients used, one would typically reach a stable oscillatory state already after about 50-100 time points). Spatio-temporal oscillations in the concentration of decondensed chromatin (which is assumed to increase with the BFA, see Main Text) are shown in Fig 4 and Supplementary Fig 10. That such a 3-species system can produce an oscillating Turing pattern under the right parameter conditions can also be seen by analyzing the reaction-diffusion graph topology as discussed in e.g.^58^.

## References

1 Meyers, M. A., Chen, P.-Y., Lin, A. Y.-M. & Seki, Y. Biological materials: Structure and mechanical properties. Progress in Materials Science 53, 1–206 (2008). https://doi.org:https://doi.org/10.1016/j.pmatsci.2007.05.002

2 Hu, S. et al. Mechanical anisotropy of adherent cells probed by a three-dimensional magnetic twisting device. Am J Physiol Cell Physiol 287, C1184–1191 (2004). https://doi.org:10.1152/ajpcell.00224.2004

3 Campas, O. et al. Quantifying cell-generated mechanical forces within living embryonic tissues. Nat Methods 11, 183–189 (2014). https://doi.org:10.1038/nmeth.2761

4 Hasnain, I. A. & Donald, A. M. Microrheological characterization of anisotropic materials. Phys Rev E Stat Nonlin Soft Matter Phys 73, 031901 (2006). https://doi.org:10.1103/PhysRevE.73.031901

5 Boon, J. P. & Yip, S. Molecular Hydrodynamics. (Dover Publications, 1991).

6 Tao, N. J., Lindsay, S. M. & Rupprecht, A. The dynamics of the DNA hydration shell at gigahertz frequencies. Biopolymers 26, 171–188 (1987). https://doi.org:10.1002/bip.360260202

7 Adichtchev, S. V. et al. Brillouin spectroscopy of biorelevant fluids in relation to viscosity and solute concentration. Phys Rev E 99, 062410 (2019). https://doi.org:10.1103/PhysRevE.99.062410

8 Bailey, M. et al. Viscoelastic properties of biopolymer hydrogels determined by Brillouin spectroscopy: A probe of tissue micromechanics. Science Advances 6, eabc1937 (2020). https://doi.org:10.1126/sciadv.abc1937

9 Yan, G., Monnier, S., Mouelhi, M. & Dehoux, T. Probing molecular crowding in compressed tissues with Brillouin light scattering. Proceedings of the National Academy of Sciences 119, e2113614119 (2022). https://doi.org:doi:10.1073/pnas.2113614119

10 Hyman, A. A., Weber, C. A. & Julicher, F. Liquid-liquid phase separation in biology. Annu Rev Cell Dev Biol 30, 39–58 (2014). https://doi.org:10.1146/annurev-cellbio-100913-013325

11 Berne, B. J. & Pecora, R. Dynamic Light Scattering: With Applications to Chemistry, Biology, and Physics. (Dover Publications, 2000).

12 Antonacci, G. et al. Recent progress and current opinions in Brillouin microscopy for life science applications. Biophysical Reviews (2020). https://doi.org:10.1007/s12551-020-00701-9

13 Palombo, F. et al. Biomechanics of fibrous proteins of the extracellular matrix studied by Brillouin scattering. J R Soc Interface 11, 20140739 (2014). https://doi.org:10.1098/rsif.2014.0739

14 Koski, K. J., Akhenblit, P., McKiernan, K. & Yarger, J. L. Non-invasive determination of the complete elastic moduli of spider silks. Nat Mater 12, 262–267 (2013). https://doi.org:10.1038/nmat3549

15 Eltony, A. M., Shao, P. & Yun, S.-H. Measuring mechanical anisotropy of the cornea with Brillouin microscopy. Nature Communications 13, 1354 (2022). https://doi.org:10.1038/s41467-022-29038-5

16 Scarcelli, G. et al. Noncontact three-dimensional mapping of intracellular hydromechanical properties by Brillouin microscopy. Nat Methods 12, 1132–1134 (2015). https://doi.org:10.1038/nmeth.3616

17 Verdonk, E. D., Wickline, S. A. & Miller, J. G. Anisotropy of ultrasonic velocity and elastic properties in normal human myocardium. The Journal of the Acoustical Society of America 92, 3039–3050 (1992). https://doi.org:10.1121/1.404200

18. Crank, J. The mathematics of diffusion. 2d edn, (Clarendon Press, 1975).

19 Cosgrove, D. J. Growth of the plant cell wall. Nat Rev Mol Cell Biol 6, 850–861 (2005). https://doi.org:10.1038/nrm1746

20 Zhang, Y. et al. Molecular insights into the complex mechanics of plant epidermal cell walls. Science 372, 706–711 (2021). https://doi.org:doi:10.1126/science.abf2824

21 Xi, X., Kim, S. H. & Tittmann, B. Atomic force microscopy based nanoindentation study of onion abaxial epidermis walls in aqueous environment. Journal of Applied Physics 117, 024703 (2015). https://doi.org:10.1063/1.4906094

22 Wang, X., Wilson, L. & Cosgrove, D. J. Pectin methylesterase selectively softens the onion epidermal wall yet reduces acid-induced creep. Journal of Experimental Botany 71, 2629–2640 (2020). https://doi.org:10.1093/jxb/eraa059

23 Haas, K. T., Wightman, R., Meyerowitz, E. M. & Peaucelle, A. Pectin homogalacturonan nanofilament expansion drives morphogenesis in plant epidermal cells. Science 367, 1003–1007 (2020). https://doi.org:10.1126/science.aaz5103

24 Trinh, D.-C. et al. How Mechanical Forces Shape Plant Organs. Current Biology 31, R143–R159 (2021). https://doi.org:https://doi.org/10.1016/j.cub.2020.12.001

25 Gadalla, A., Dehoux, T. & Audoin, B. Transverse mechanical properties of cell walls of single living plant cells probed by laser-generated acoustic waves. Planta 239, 1129–1137 (2014).

26 Elsayad, K. et al. Mapping the subcellular mechanical properties of live cells in tissues with fluorescence emission-Brillouin imaging. Sci Signal 9, rs5 (2016). https://doi.org:10.1126/scisignal.aaf6326

27. Bacete, L. et al. THESEUS1 modulates cell wall stiffness and abscisic acid production in

28. Arabidopsis thaliana. Proceedings of the National Academy of Sciences 119, e2119258119 (2022). https://doi.org:doi:10.1073/pnas.2119258119

28 Monshausen, G. B., Messerli, M. A. & Gilroy, S. Imaging of the Yellow Cameleon 3.6 Indicator Reveals That Elevations in Cytosolic Ca2+ Follow Oscillating Increases in Growth in Root Hairs of Arabidopsis Plant Physiology 147, 1690-1698 (2008). https://doi.org:10.1104/pp.108.123638

29 Rygol, J., Büchner, K.-H., Winter, K. & Zimmermann, U. Day/night variations in turgor pressure in individual cells of Mesembryanthemum crystallinum L. Oecologia 69, 171–175 (1986). https://doi.org:10.1007/BF00377617

30 Ou, H. D. et al. ChromEMT: Visualizing 3D chromatin structure and compaction in interphase and mitotic cells. Science 357, eaag0025 (2017). https://doi.org:10.1126/science.aag0025

31 Hansen, J. C., Maeshima, K. & Hendzel, M. J. in Epigenetics and Chromatin Vol. 14 (BioMed Central Ltd, 2021).

32 Gibson, B. A. et al. In diverse conditions, intrinsic chromatin condensates have liquid-like material properties. Proc Natl Acad Sci U S A 120, e2218085120 (2023). https://doi.org:10.1073/pnas.2218085120

33 Matsushita, K. et al. Intranuclear mesoscale viscoelastic changes during osteoblastic differentiation of human mesenchymal stem cells. The FASEB Journal 35, e22071 (2021). https://doi.org:https://doi.org/10.1096/fj.202100536RR

34 Yesbolatova, A. K., Arai, R., Sakaue, T. & Kimura, A. Formulation of Chromatin Mobility as a Function of Nuclear Size during C. elegans Embryogenesis Using Polymer Physics Theories. Physical Review Letters 128, 178101 (2022). https://doi.org:10.1103/PhysRevLett.128.178101

35 Khanna, N., Zhang, Y., Lucas, J. S., Dudko, O. K. & Murre, C. Chromosome dynamics near the sol-gel phase transition dictate the timing of remote genomic interactions. Nature Communications 10, 2771 (2019). https://doi.org:10.1038/s41467-019-10628-9

36 Eshghi, I., Eaton, J. A. & Zidovska, A. Interphase Chromatin Undergoes a Local Sol-Gel Transition upon Cell Differentiation. Physical Review Letters 126, 228101 (2021). https://doi.org:10.1103/PhysRevLett.126.228101

37 Zidovska, A. The rich inner life of the cell nucleus: dynamic organization, active flows, and emergent rheology. Biophysical Reviews 12, 1093–1106 (2020). https://doi.org:10.1007/s12551-020-00761-x

38 Zhang, J. et al. Nuclear Mechanics within Intact Cells Is Regulated by Cytoskeletal Network and Internal Nanostructures. Small 16, 1907688 (2020). https://doi.org:https://doi.org/10.1002/smll.201907688

39 Schlüßler, R. et al. Correlative all-optical quantification of mass density and mechanics of subcellular compartments with fluorescence specificity. eLife 11, e68490 (2022). https://doi.org:10.7554/eLife.68490

40 Fullgrabe, J., Hajji, N. & Joseph, B. Cracking the death code: apoptosis-related histone modifications. Cell Death Differ 17, 1238–1243 (2010). https://doi.org:10.1038/cdd.2010.58

41 Cui, Y. & Bustamante, C. Pulling a single chromatin fiber reveals the forces that maintain its higher-order structure. Proc Natl Acad Sci U S A 97, 127–132 (2000). https://doi.org:10.1073/pnas.97.1.127

42 Ghavanloo, E. Persistence length of collagen molecules based on nonlocal viscoelastic model. J Biol Phys 43, 525–534 (2017). https://doi.org:10.1007/s10867-017-9467-2

43 Chaikin, P. M. & Lubensky, T. C. Principles of Condensed Matter Physics. (Cambridge University Press, 1995).

44 Strick, R., Strissel, P. L., Gavrilov, K. & Levi-Setti, R. Cation–chromatin binding as shown by ion microscopy is essential for the structural integrity of chromosomes. Journal of Cell Biology 155, 899–910 (2001). https://doi.org:10.1083/jcb.200105026

45 Lebeaupin, T., Smith, R. & Huet, S. in Nuclear Architecture and Dynamics Vol. 2 (eds Christophe Lavelle & Jean-Marc Victor) 209-232 (Academic Press, 2018).

46 Gerace, L. Molecular trafficking across the nuclear pore complex. Current Opinion in Cell Biology 4, 637–645 (1992). https://doi.org:https://doi.org/10.1016/0955-0674(92)90083-O

47 Remer, I., Shaashoua, R., Shemesh, N., Ben-Zvi, A. & Bilenca, A. Publisher Correction: High-sensitivity and high-specificity biomechanical imaging by stimulated Brillouin scattering microscopy. Nat Methods 17, 1060 (2020). https://doi.org:10.1038/s41592-020-0956-z

48 Anderson, P. W. More Is Different. Science 177, 393–396 (1972). https://doi.org:10.1126/science.177.4047.393

49 Lee, C. F. & Wurtz, J. D. Novel physics arising from phase transitions in biology. Journal of Physics D: Applied Physics 52, 023001 (2018). https://doi.org:10.1088/1361-6463/aae510

50 Gompper, G. et al. The 2020 motile active matter roadmap. Journal of Physics: Condensed Matter 32, 193001 (2020). https://doi.org:10.1088/1361-648X/ab6348

51 Schreiber, B., Elsayad, K. & Heinze, K. G. Axicon-based Bessel beams for flat-field illumination in total internal reflection fluorescence microscopy. Opt Lett 42, 3880–3883 (2017). https://doi.org:10.1364/OL.42.003880

52 Shijun, X., Weiner, A. M. & Lin, C. A dispersion law for virtually imaged phased-array spectral dispersers based on paraxial wave theory. IEEE Journal of Quantum Electronics 40, 420–426 (2004). https://doi.org:10.1109/JQE.2004.825210

53 Edrei, E., Gather, M. C. & Scarcelli, G. Integration of spectral coronagraphy within VIPA-based spectrometers for high extinction Brillouin imaging. Opt Express 25, 6895–6903 (2017). https://doi.org:10.1364/OE.25.006895

54 Murashige, T. & Skoog, F. A Revised Medium for Rapid Growth and Bio Assays with Tobacco Tissue Cultures. Physiologia Plantarum 15, 473–497 (1962). https://doi.org:https://doi.org/10.1111/j.1399-3054.1962.tb08052.x

55 Peaucelle, A., Wightman, R. & Höfte, H. The Control of Growth Symmetry Breaking in the Arabidopsis Hypocotyl. Current Biology 25, 1746–1752 (2015). https://doi.org:10.1016/j.cub.2015.05.022

56 Elsayad, K. et al. Mechanical Properties of cellulose fibers measured by Brillouin spectroscopy. Cellulose (2020). https://doi.org:10.1007/s10570-020-03075-z

57 Wang, S. et al. Biomechanical assessment of myocardial infarction using optical coherence elastography. Biomed Opt Express 9, 728–742 (2018). https://doi.org:10.1364/BOE.9.000728

58 Diego, X., Marcon, L., Müller, P. & Sharpe, J. Key Features of Turing Systems are Determined Purely by Network Topology. Physical Review X 8, 021071 (2018). https://doi.org:10.1103/PhysRevX.8.021071

