## Supplementary Material for "Imaging the microscopic viscoelastic anisotropy in living cells"

#### Supplementary Figures:

**Supplementary Figure 1:** Concept and implementation of radial-VIPA for BLS microspectroscopy.

**Supplementary Figure 2:** Alignment correction and spatial resolution of radial-VIPA based microspectroscopy setup.

**Supplementary Figure 3:** Anisotropy of muscle myofiber during drying induced hypercontraction.

**Supplementary Figure 4:** Anisotropy in *Arabidopsis* cell wall elastic moduli.

**Supplementary Figure 5:** Viscoelastic anisotropy maps of anticlinal cell walls of *Arabidopsis thaliana* and effect of anisotropy on the dispersion of *M'* and *M''*.

**Supplementary Figure 6:** Brillouin Fractional Anisotropy (BFA) in the nucleus during apoptosis and treatment.

**Supplementary Figure 7:** Correlation of BFA to SIR-DNA fluorescence and to nuclear morphology in U2OS cells.

**Supplementary Figure 8:** Scaling of the orientational correlation function.

**Supplementary Figure 9:** Scaling of elastic storage moduli in mixed Reuss-Voigt material.

**Supplementary Figure 10:** Reaction-diffusion simulations in the presence of noise and perturbed activation/inhibition rates.

#### Supplementary text

1. Radial- VIPA and Apodization filters
2. Obtaining elastic moduli from $\nu_B$
3. Spatial, temporal and spectral resolution
4. Effective Medium modeling
5. Association of Reuss *vs.* Voigt scaling with ionic concentration
6. 3-species reaction diffusion model
7. Supplementary references

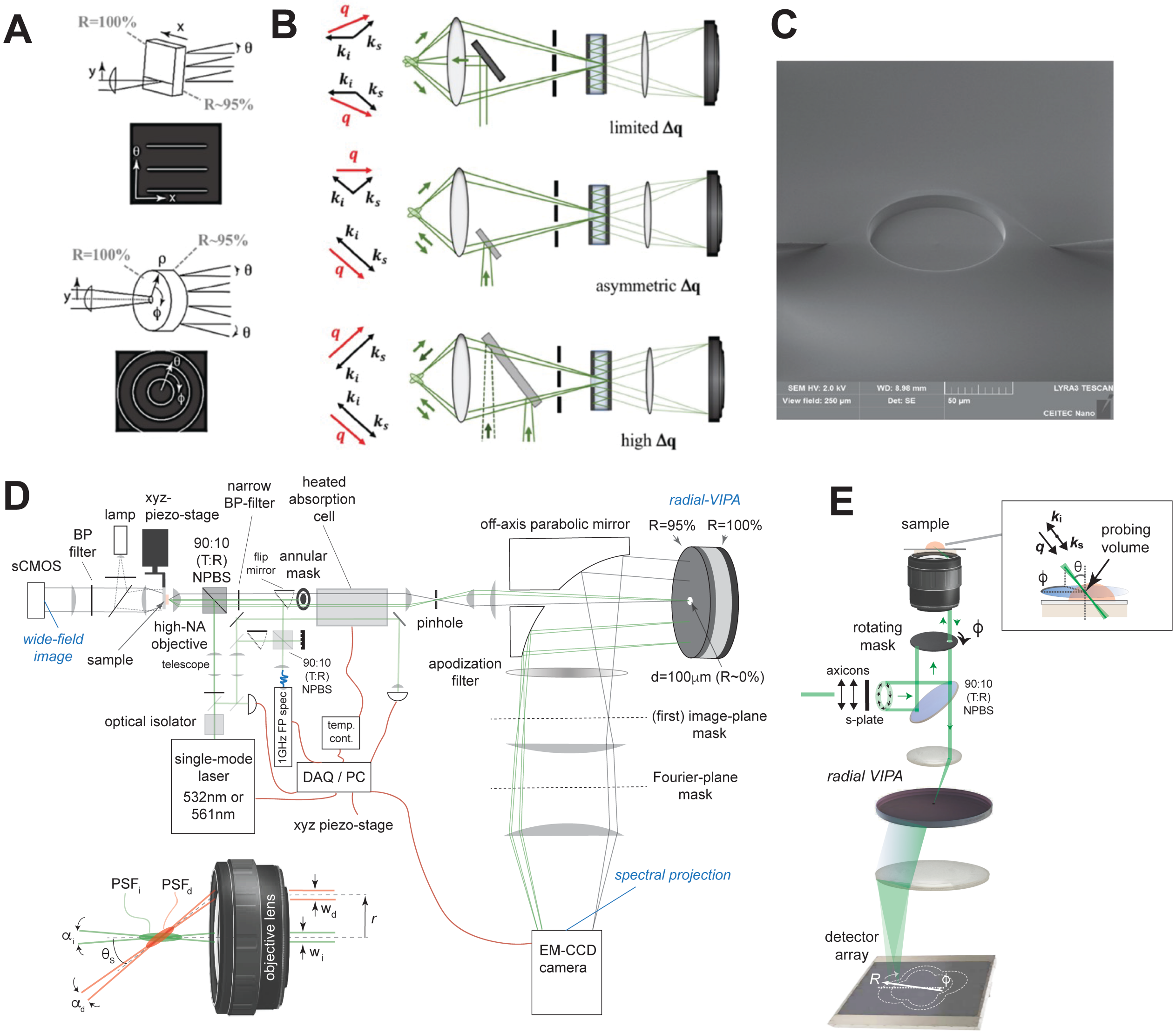

**Supplementary Fig 1: Concept and implementation of radial-VIPA for BLS microspectroscopy.** (**A**) Sketch of conventional VIPA, and radial-VIPA, showing angular spectra that would be obtained from a single frequency signal. (R=reflectivity of indicated surface). (**B**) Three different experimental configurations employing the radial-VIPA to measure angle (ϕ) -resolved BLS spectra at a given position in a sample. Also shown are the range of wavevectors ***q*** that can be probed ($\boldsymbol{k}_i$ and $\boldsymbol{k}_s$ are the incident and scattering wavevectors). (**C**) Scanning Electron Microscopy (SEM) image of 100μm diameter *entrance hole* milled in a dielectric coated etalon (radial-VIPA) using a dual-beam Focused Ion Beam (FIB)/SEM. Also visible are the two closed-loop nanomanipulator needles that were kept in contact with the substrate surface to suppress ion beam drift during the milling process. (**D**) Experimental setup for single-shot measurements of BLS anisotropy, corresponding to first design depicted in (B). (BP= Band Pass, NPBS = Non-Polarizing Beam Splitter, DAQ=Data Acquisition, FP=Fabry Perot). *Inset:* close-up of the incident and detection angles and excitation and detection Point Spread Functions ($PSF_i$ and $PSF_d$)--see supplementary text. (**E**) Schematic of an implementation of the radial-VIPA corresponding to third design depicted (B) (see supplementary text).

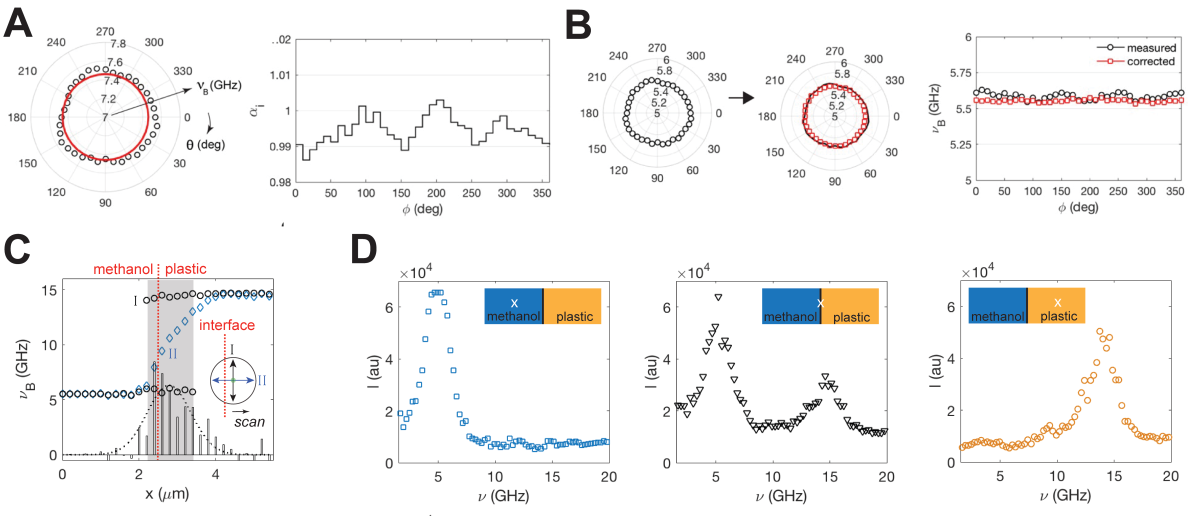


**Supplementary Fig 2: Alignment correction and spatial resolution of radial-VIPA based microspectroscopy setup. (A)** Alignment correction for radial-VIPA. *Black circles:* Example of unprocessed BLS-peak position, $\nu_B(\phi)$, of distilled water at room temperature measured with radial VIPA. *Red line:* expected value, $\nu_B^{(t)} = 7.46$ GHz. Also shown is the correction coefficient $\alpha(\phi_i) = \nu_B(\phi_i)/\nu_B^{(t)}$ for data (Methods). (**B**) Subsequent measurement (on methanol) in same imaging session showing measured $\nu_B(\phi_i)$ (*black circles* and *line*), and corrected BLS frequency shift $= \nu_B(\phi_i)/\alpha(\phi_i)$ (*red squares*). Also shown is plot of the measured and corrected values. **(C)** Lateral spatial resolution of radial-VIPA microspectroscopy setup obtained obtained from scanning across a plastic-methanol interface using the first configuration of Supplementary Fig 1B. Shown are the BLS peak positions as a function of lateral distance at azimuthal angles parallel (I) and perpendicular (II) to the interface (*inset*). The former shows two distinct peaks (corresponding roughly to the two materials), attributed to the two phonon modes in the effective PSF, localized in the two materials respectively (see also (D)). The distance *x* over which both peaks are observed corresponds to ≈ 1.5 μm. the latter shows a smooth transition between the two BLS frequency shifts of the materials, we attribute to a phonon mode that exist in both materials. The slope of the transition of $\nu_B$ with respect to *x* (in GHz/μm) is shown as a bar chart. A Gaussian fit yields $\sigma = 0.570$ μm corresponding to a FWHM of 1.34μm, suggesting a lateral spatial resolution of ≈ 1.5 μm. All analysis was done after aberration corrections in (B) using measurements of methanol away from the interface. (**D**) Brillouin Frequency shift (BFS) at three positions in methanol, near the interface and in plastic at an azimuthal angle parallel to the interface.

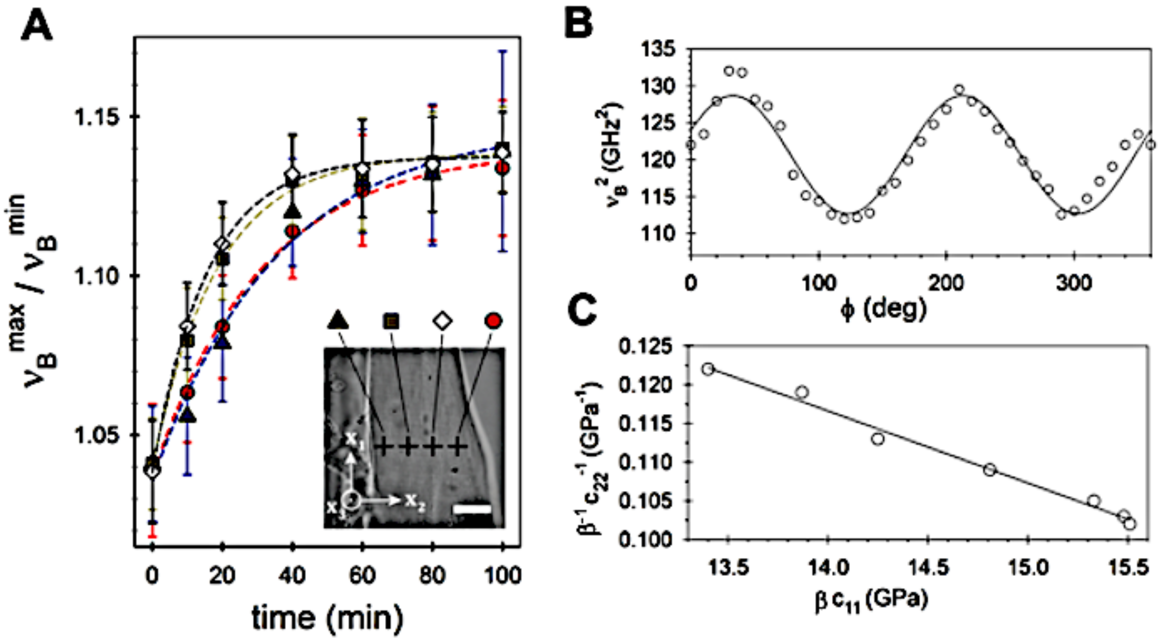


**Supplementary Fig 3: Anisotropy of muscle myofiber during drying induced hypercontraction.** (**A**) Ratio of maximum ($\nu_B^{\max}$) to minimum ($\nu_B^{\min}$) Brillouin frequency shift at different positions in a myofiber during dehydration. The anisotropy, as defined by $\nu_B^{\max}/\nu_B^{\min}$, saturates more rapidly in the inner (squares and diamonds) versus the outer (triangles and circles) positions, likely due to migration of water towards the surface where it evaporates. The decrease in the anisotropy is accompanied by an increased mean (angle averaged) value of $\nu_B = < \nu_B(\phi) >_\phi$, consistent with an increase in the contribution from the solid fraction. (**B**) Characteristic plot of $\nu_B^2(\phi)$ fitted with transverse anisotropic model (*solid line*). (**C**) Plot of scaling of the perpendicular-to-fiber-axis inverse elastic modulus *vs.* the parallel-to-fiber-axis elastic modulus at different stages of dehydration. The dimensionless coefficient $\beta \sim O(1)$, will depend on the refractive index and density, with the former assumed to a first approximation to be isotropic. Solid line shows least-square-fit for a transverse isotropic model. The data suggests a linear relationship between $c_{22}$ and $c_{11}^{-1}$ suggesting a transverse isotropic elastic symmetry is maintained during dehydration (supplementary text).

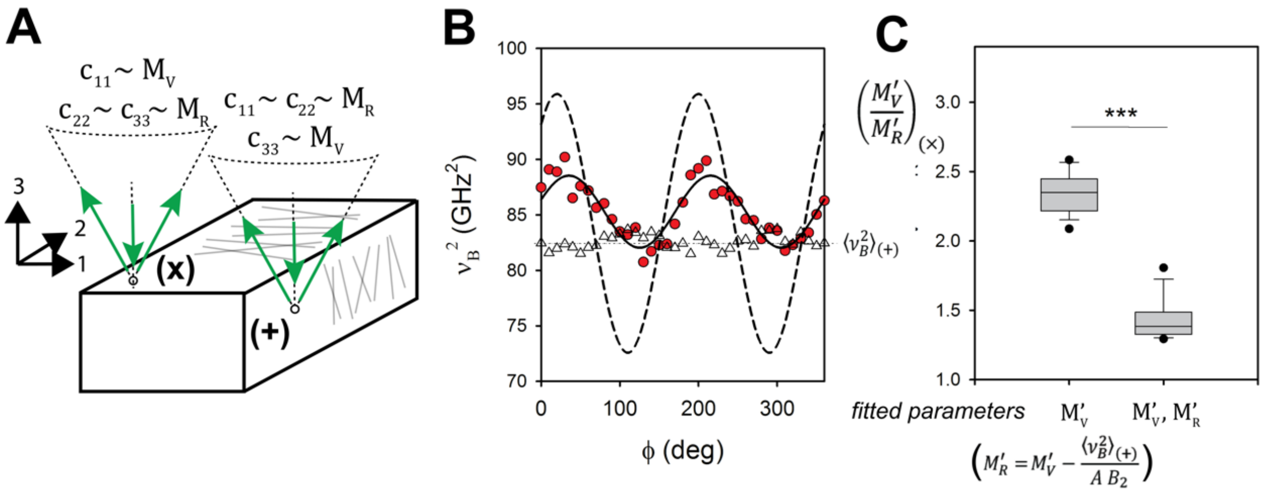


**Supplementary Fig 4: Anisotropy in *Arabidopsis* cell wall elastic moduli.** (**A**) Schematic showing effective medium approximations of the three diagonal stiffness tensor components $(c_{11}, c_{22}, c_{33})$ employed. $M_V$ =Voigt averaging, $M_R$ =Reuss averaging. See Supplementary Text. (**B**) $\nu_B^2(\phi)$ for two cell walls of the same cell. *Red circles* = outer cell wall ("x" in Fig. S4A), *white triangles* = inner cell wall ("+" in Fig. S4A). Solid black line shows fit of the former with Eqn S9A in Supplementary Text. Dashed black line shows fit of the same data but with constraint imposed between $M'_V$ and $M'_R$ from fitting of the inner cell wall (Eqn S9B in Supplementary Text). This suggests inner ("+") and outer ("x") walls have distinct BLS elastic anisotropy. (**C**) Difference in the ratio $M'_V/M'_R$ of outer cell walls obtained with and without the constraint set from fitting the inner cell walls (N=15 cells). This suggests that the effective longitudinal modulus parallel to the microfibrils ($M'_V$) relative to that perpendicular ($M'_R$) is smaller in the outer walls ("x") than the inner walls ("+"); namely that the outer walls are more isotropic than the inner walls.

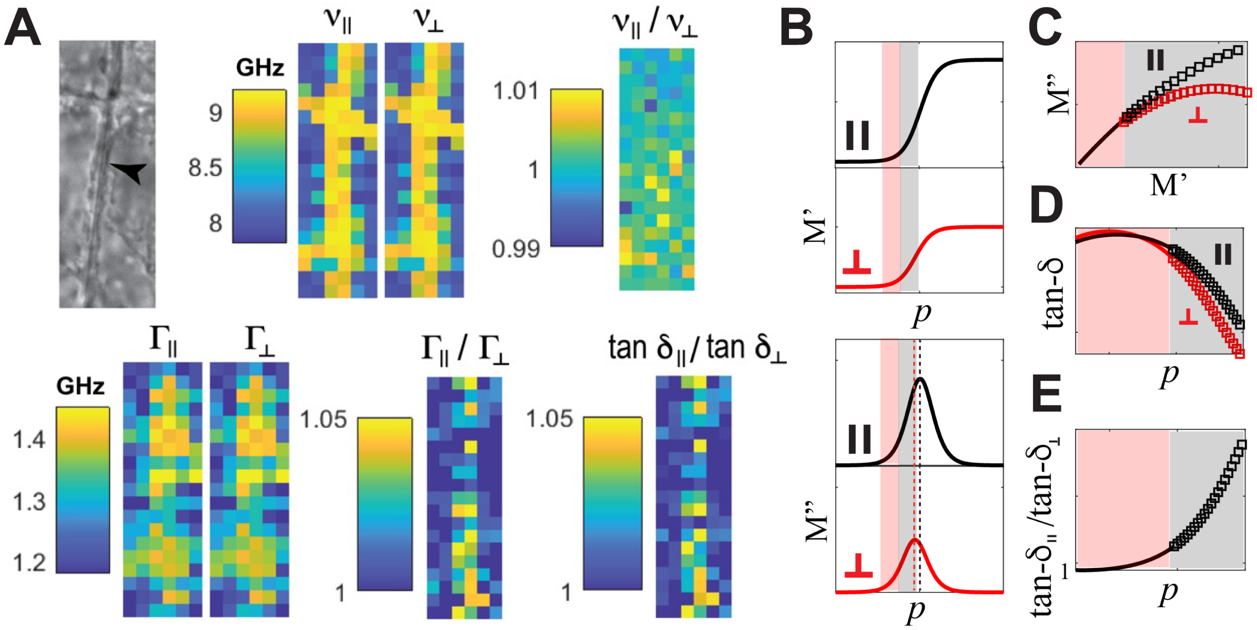


**Supplementary Fig 5: Viscoelastic anisotropy maps of anticlinal walls of *Arabidopsis thaliana* hypocotyl cells, and effect of anisotropy on the dispersion of M' and M''. (A)** Maps of the Brillouin frequency shift $\nu_B$, and line-width $\Gamma_B$ parallel (||) and perpendicular (⊥) to the growth axis (see schematic in inset of Fig 2D) and the ratios thereof. Also shown is the ratio of the loss tangents ($\tan\delta$). **(B)** Longitudinal storage M' and Loss M'' moduli as a function of a parameter that embodies the degree of methylation (*p*) for distinct relaxation times parallel (||) and perpendicular (⊥) to the cell growth axis. If for the *pmeo5* mutants *p* only takes on smaller values (shaded red), this would result in both a lower storage modulus and more isotropic mechanical behavior that is observed [(**C**)-(**E**)]. The predicted functional scaling of *M'* vs. *M''* shown in (**C**) can be seen to be similar to that experimentally observed in Fig 2F ($\tan\delta = M''/M'$).

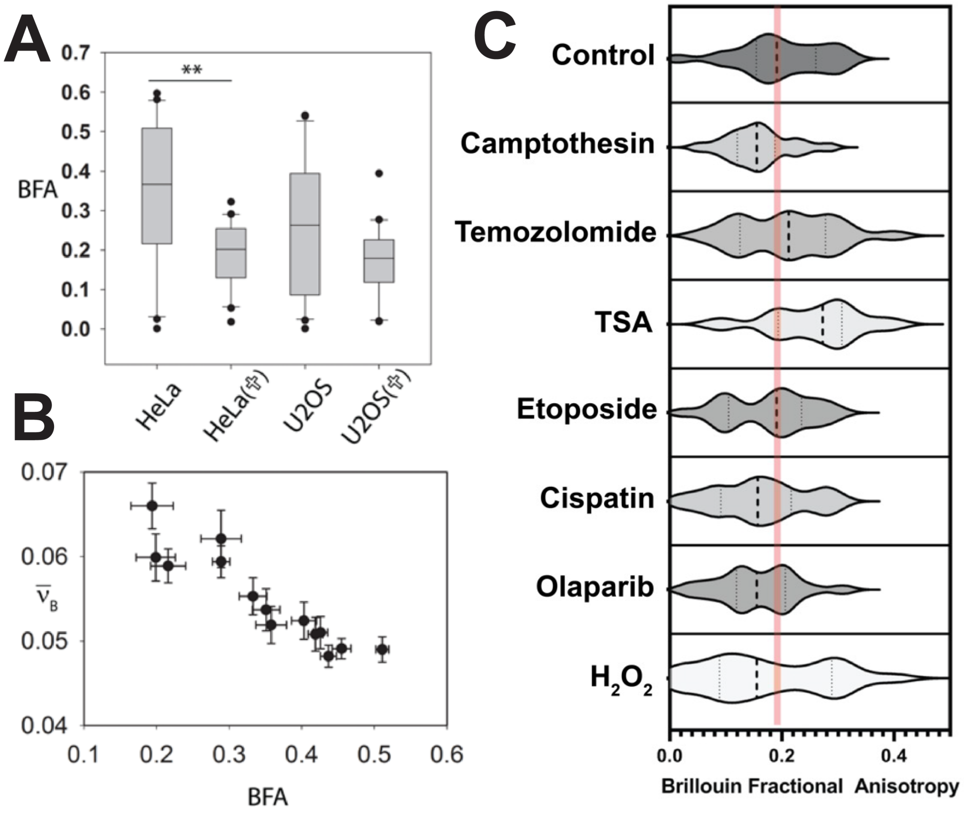


**Supplementary Fig 6: Brillouin Fractional Anisotropy (BFA) in the nucleus during apoptosis and treatment.** (**A**) Mean BFA in the nucleus of healthy interface HeLa and U2OS cells and cells during apoptosis (✞)**.** Based on a Mann-Whitney test the decrease in the value of the BFA is significant (P = 0.005, Mann–Whitney U =95, n=20) for the case of HeLa cells, but not for the case of U2OS cells (P =0.189 , Mann–Whitney U =151, n =20). An Ansari-Bradley test performed on median subtracted data suggests a significant difference in the distributions of the BFA during apoptosis for both cell types. **(B)** Mean value of angle-average Brillouin elastic contrast ($\overline{\nu_B}$) (see Supplementary Text) over entire nuclei as a function of nuclear averaged BFA for 15 interphase HeLa cells, showing an inverse correlation between the two (P<0.0001, Pearson correlation coefficient $\rho$ =- 0.918). **(C)** Brillouin Fractional Anisotropy BFA for cells subject to various treatments. Red line indicates median value for untreated cells. U2OS cells were treated with 25 µM camptothecin for 1h, 1 mM temozolomide for 4h, 10 µM trichostatin A (TSA) for 6h, 20 µM etoposide for 1h, 30 µM cisplatin for 2h, 10 µM olaparib for 2h, and 200 µM $H_2O_2$ for 10min.

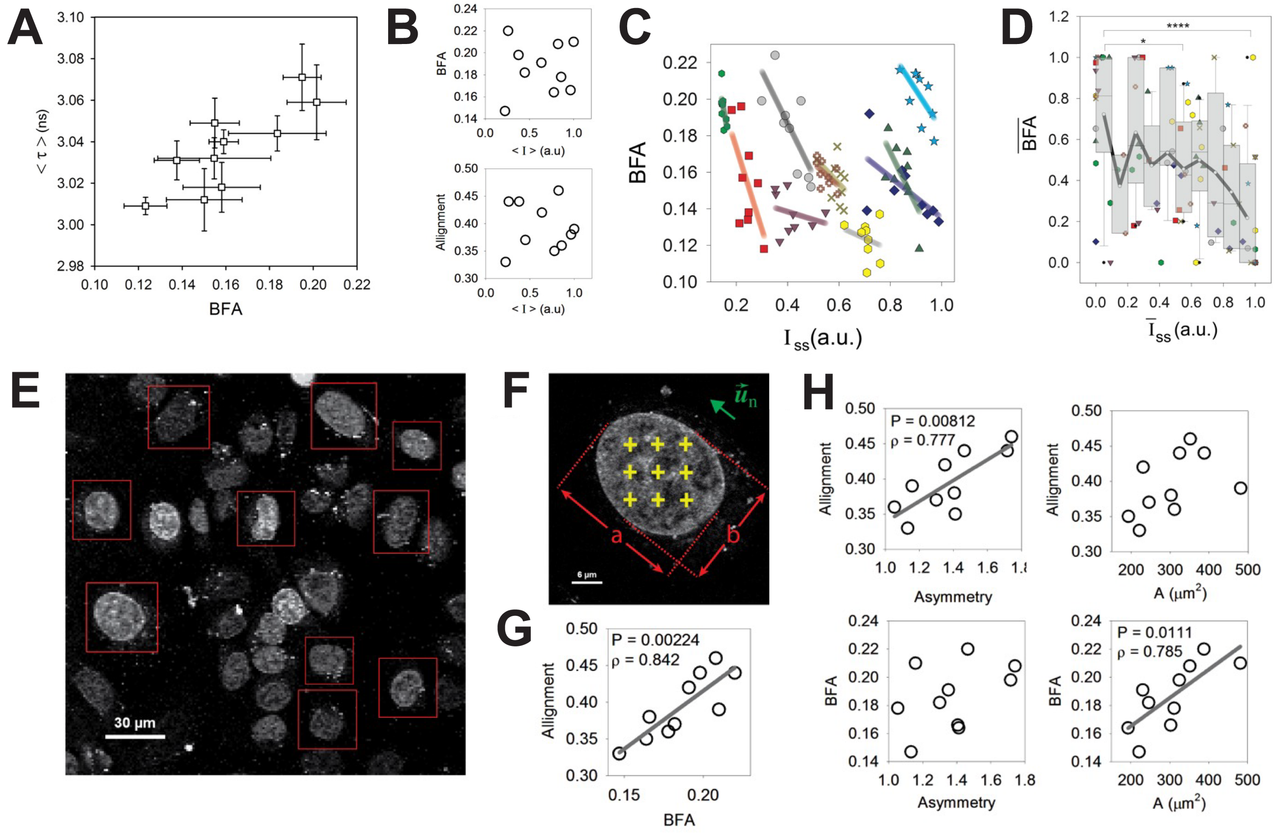

**Supplementary Fig 7: Correlation of BFA to SIR-DNA fluorescence and to nuclear morphology in U2OS cells. (A)** Average SIR-DNA fluorescence lifetime in relation to nucleus-averaged BFA for 10 U2OS cells, showing a positive correlation (P=0.0036, Pearson correlation coefficient $\rho$ = 0.821). **(B)** Nucleus average steady-state fluorescence intensity (<I>) as a function of nucleus-averaged BFA and alignment parameter (see Methods) suggesting no clear correlation. **(C)** Relation between local-BFA and steady-state fluorescence intensity for 10 cells, show a negative correlation on an individual cell case. Different symbols correspond to different cells, and lines are least-square linear fits for data from each cell. **(D)** Normalized fluorescence intensity and BFA for each of the cells in (C), calculated as $\overline{BFA} = [BFA - \min(BFA)]/[\max(BFA) - \min\,(BFA)]$ and $\bar{I} = [I - \min(I)]/[\max(I) - \min\,(I)]$. Solid black line shows mean value for binned data ($\Delta\bar{I} = 0.1$) suggesting a negative correlation (**E**) Confocal fluorescence image of SIR-DNA stained U2OS cells. (**F**) Characteristic individual nucleus confocal fluorescence image with points (yellow crosses) at which BLS-grid is scanned. (**G**) *Alignment* defined as $|\vec{\boldsymbol{u}}_{\boldsymbol{n}} \cdot \vec{\boldsymbol{u}}|$ where $\vec{\boldsymbol{u}}$ is the unit-direction of the average BLS anisotropy (Methods) and $\vec{\boldsymbol{u}}_{\boldsymbol{n}}$ the direction of the long-axis of nucleus, indicated in (B), as a function of the nucleus-averaged BFA obtained from grid scans (N=10 cells), showing a positive correlation. (**H**) Dependence of *Alignment* and nucleus-averaged BFA on the nuclear *asymmetry* (calculated as $a/b$, where $a$ and $b$ are as defined in (B)), and nuclear area $A$. There is a positive correlation between the nucleus-averaged BFA and the nuclear area (P=0.00111, Pearson correlation coefficient $\rho$ = 0.785), as well as between the *alignment* and both the BFA (P=0.00224, $\rho$ = 0.842) and nuclear asymmetry (P=0.00812, $\rho$ = 0.777). This suggests the nuclear-averaged internal BLS elastic anisotropy (as defined by the BFA) is preferentially aligned with the nuclear symmetry axis and that this is more pronounced for larger nuclei.

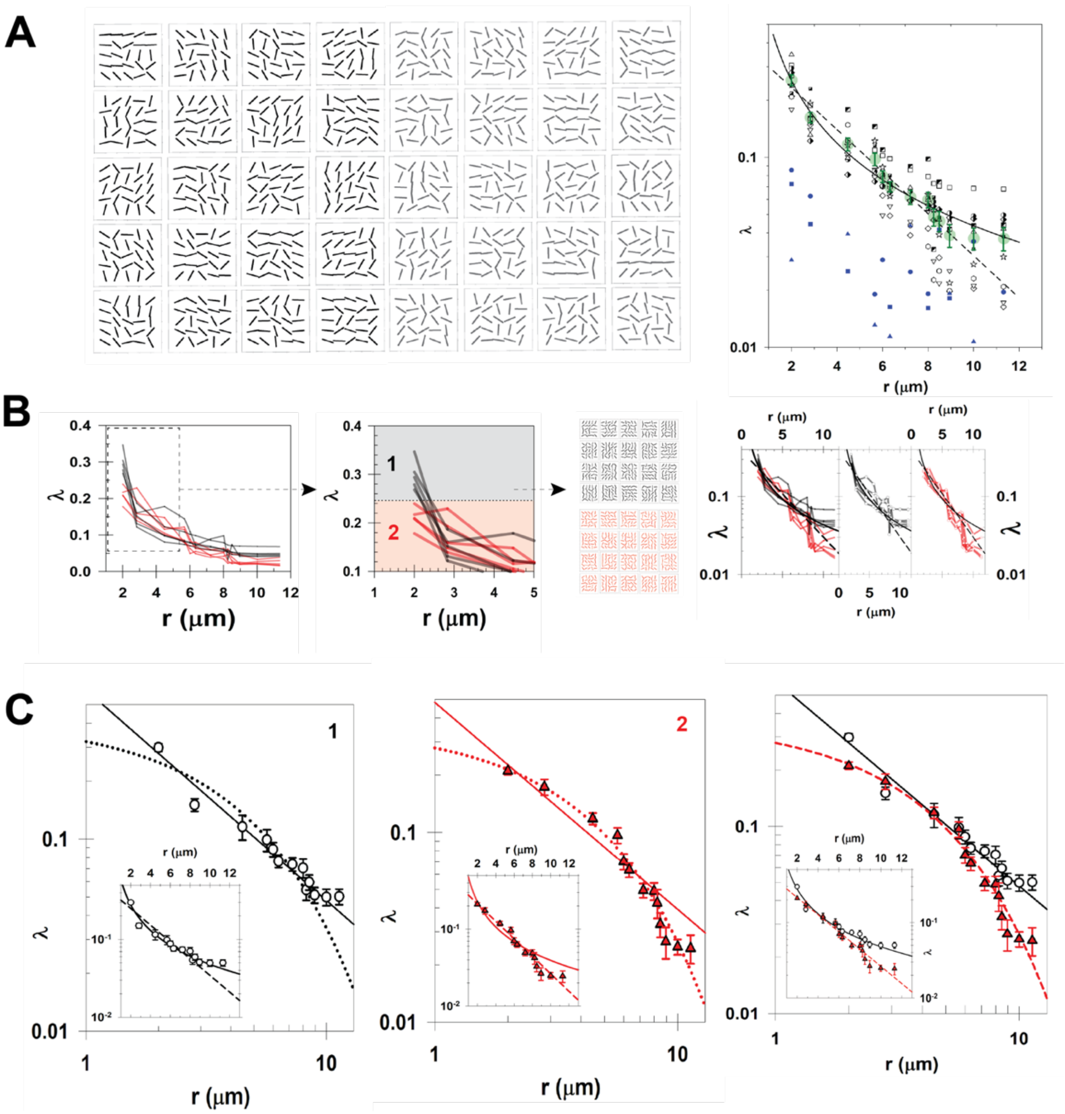


**Supplementary Fig 8: Scaling of the orientational correlation function.** (**A**) Examples of nuclear rapid scan data (2μm grid-step size) used to calculate angular correlation functions (λ) (Methods). Also shown is plot of the calculated $\lambda$ for individual cells (open symbols and half-shaded symbols) and mean $\bar{\lambda}$ (large solid green circles) for non-apoptotic cells, as a function of distance plotted on a semilogarithmic axis. Blue symbols are apoptotic cells. $\bar{\lambda}(r)$ can be described equally well by a two-parameter exponential and power law function (dashed and solid lines show respective least squares fits). By dividing cell data into two subgroups composed of cells with a higher and lower $\lambda$ at short distances $r$ [we set this cut-off at $\lambda(r$=2μm)=0.25] we observe two distinct scaling behaviors, as shown as *red* and *black* lines in (**B**). The mean $\bar{\lambda}$ for these two subgroups is plotted in (**C**). *Open black symbols* = $\lambda(r$=2μm)>0.25, and *solid red symbols* = $\lambda(r$=2μm)<0.25). The former [$\lambda(r$=2μm)>0.25] is seen to be better described by a power law than an exponential function (adjusted $R^2$=0.95 versus 0.86), whereas the latter [$\lambda(r$=2μm)<0.25] are better described by an exponential than a power law function (adjusted $R^2$=0.99 versus 0.93). Solid lines represent power law fits, whereas dashed and dotted lines represent exponential fits. The effective length scale (distance $r$ for $\lambda(r)$ to drop by 1/e) is 4.04μm for the subset of cells with $\lambda(r$=2μm)>0.25, compared to 3.82μm for those with $\lambda(r$=2μm)<0.25.

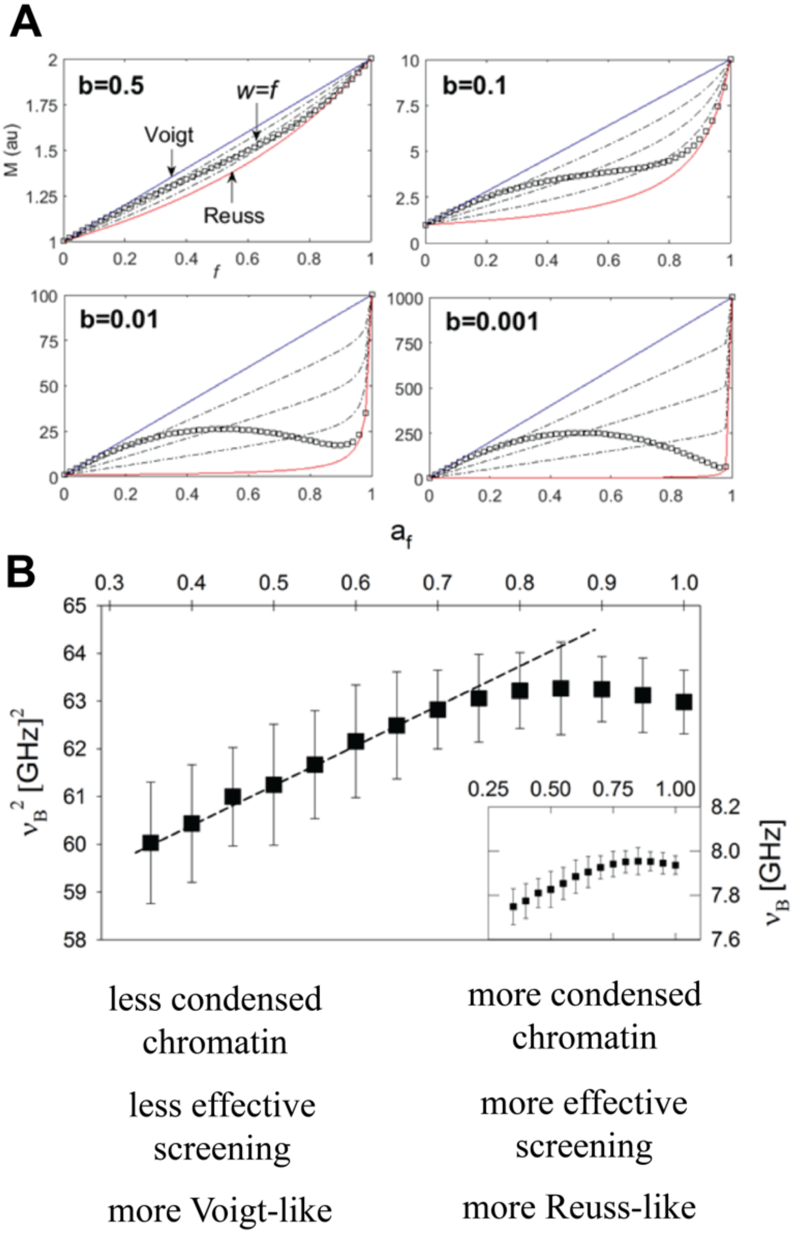


**Supplementary Fig 9: Scaling of elastic storage moduli in mixed Reuss-Voigt material.** (**A**) Elastic storage modulus as a function of filling factor $f$. Blue and red lines show pure Voigt and Reuss models viz. $w = 1$ and $w = 0$ (see Eqns 6 & 7, Methods). Squares show case for $w = f$ (*i.e.* contribution of Reuss relative to Voigt scaling increases with filling factor). The parameter $b$ is the ratio of the liquid to solid moduli (Eqns 7). (**B**) Measured BLS frequency shift squared ($\nu_B^2 \propto M$) in interphase cell nuclei as a function of normalized SIR-DNA fluorescence intensity ($a_f$)—which can be considered to scale with degree of condensed chromatin (Fig S6 and S7). The observed increase followed by a saturation and slight decrease in $\nu_B^2$ for increasing $a_f$ is consistent with a transition between Voigt and Reuss-like scaling (coupled to decoupled chromatin-solvent dynamics) as can be expected to occur for increasing cation concentration and screening.

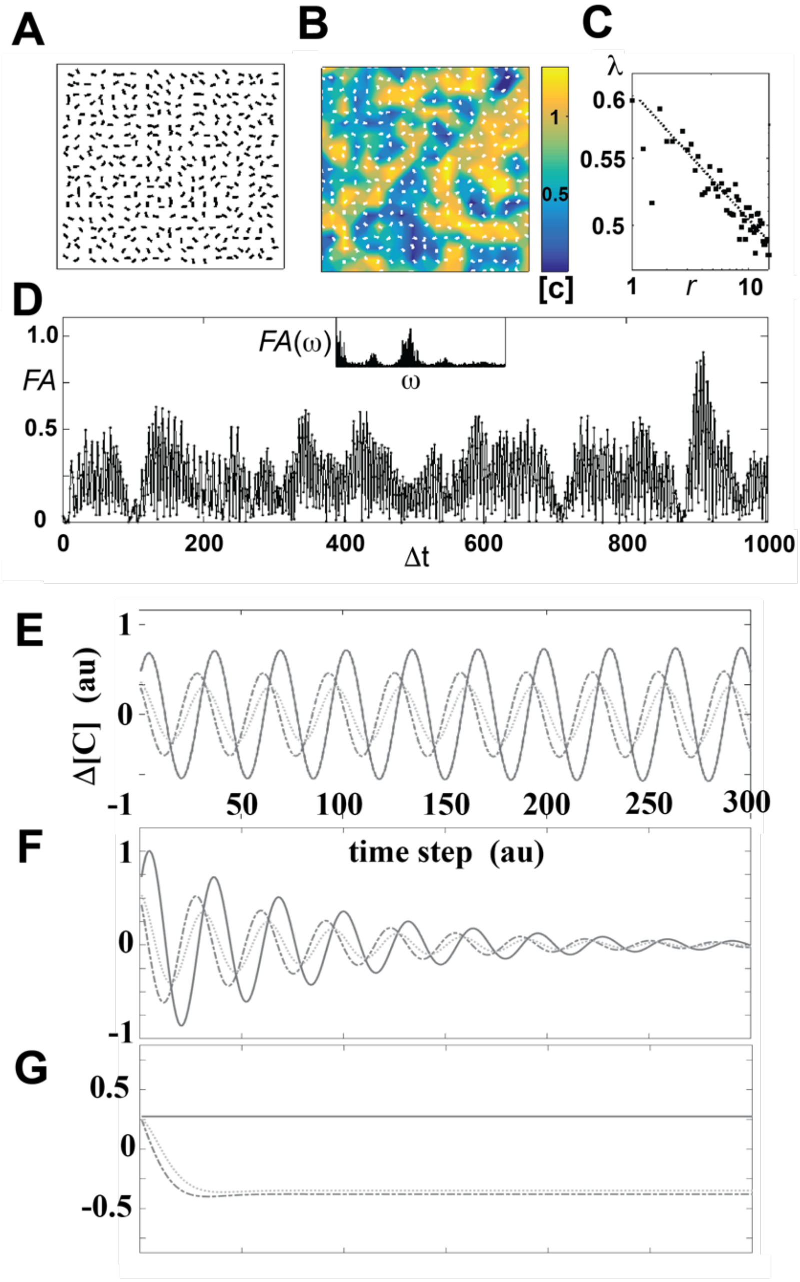


**Supplementary Fig 10: Reaction-diffusion model simulations in the presence of noise and perturbed activation/inhibition rates.** Simulations for parameters that give a spatio-temporally oscillatory (STO) solution in the presence of 2% white noise (in concentration of each species) with a parabolic restoring force $\propto \left(k_{ij} - k_{ij}^{c}\right)^2$. Shown are (**A**) the calculated local directors $\boldsymbol{u}$, (**B**) superimposed on the relative condensed chromatin concentration [C], and (**C**) the calculated order parameter λ as described in Methods and Supplementary Text. (**D**) Example time trace of [C] at a given point. (*Inset*: corresponding power spectrum). (**E**)-(**G**) show simulation results of [C], [+] and [x] at a given spatial position, (**E**) in the absence of noise, (**F**) when the activation rate k([+]→[C]) is perturbed from the STO value by 2% (**F**) and by 10% (**G**) (see also Supplementary Text). These results suggest that while oscillations may be stable to small random concentration perturbations, they are acutely sensitive to the mean values of the activation rates.

### Supplementary Text

#### 1. Radial- VIPA and Apodization filters

*Radial-VIPA fabrication:* A number of Radial VIPA's were fabricated with different specifications. For the results reported in this manuscript, 25.5 $mm^2$ diameter ultra-high flatness etalon substrates (Laseroptik GmbH, DE) with a thicknesses of 3mm were used. The reflective coatings (R=95% on one side and 100% on the other) either consisted of sputtered Au metal films (as described below) or propriety reflective dielectric coatings (fabricated by Laseroptik GmbH, DE). With the dielectric films a higher finesse was generally achievable (routinely >80) compared to metal coated films. In addition, the dielectric coated etalons also had the advantage that they would not degrade with time (whereas the metal coated ones would after several weeks). This however can possibly be in part circumvented by employing an additional coating layer.

The centric hole with Ø=100 μm was micromachined on one side of the etalon metal coatings using a TESCAN dual-beam FIB/SEM LYRA3 system under 30 kV accelerating voltage and 165 pA probe current. For the dielectric coated etalons, the same hole structure with Ø=100 μm was milled through the ~12 μm thick stack of high- and low-index silica multilayer of the etalons, which constitute the reflective layers, under 30 kV accelerating voltage and 660 pA probe current. To ensure the accuracy of the etching process upon a poorly conductive surface, two closed-loop nanomanipulator needles equipped in the LYRA3 system were continuously in contact with the substrate surface to suppress ion beam drift during the milling process. The surface profiles of the holes were inspected by Scanning Electron Microscopy (SEM)–Fig S1B, where the two nanomanipulator needles used to suppress ion beam drift are also shown. The extremely sharp boundaries (edges) help in assuring a high finesse (which is in part also dependent on the sharpness of this transition).

To realize the metal coated VIPA, the coating thickness that corresponds to 95% reflectivity was established on test samples with different thicknesses under the same fabrication conditions, from which it was confirmed that a thickness of ≈70nm was required. On one side of the etalons a 70 nm thin gold film was subsequently deposited by Ion Beam Sputtering (IBS), at 5 Å $s^{-1}$, including a 2 nm titanium adhesion layer that was deposited at 1 Å $s^{-1}$. The coatings were realized by an in-house developed sputterer equipped with a Kaufman-type argon ion source. The flow of argon gas into the deposition chamber was controlled by a flow-controller (MKS Mass Flo 1179 C with Type 247 readout) and set to 5.5 sccm. The chamber was pumped by a cryo-pump (Leybold CoolVac 2000CL), the background pressure in the chamber was 1.5 x $10^{-6}$ mbar (measured by Edwards Wide Range Gauge WRG-S-NW25), and the operating pressure was 2.2 x$10^{-5}$ mbar. Argon ions were extracted from the source and accelerated (kinetic energy 600 eV) towards a gold sputtering target (purity 99.99 %, MaTeck) by a set of molybdenum grids. The sputtering angle (angle between the axis of argon ion broad-beam and surface normal of sputtered target) was approximately 35° with respect to a gold target surface normal. A holder with substrates was oriented parallel to the sputtered target. The thickness of deposited film was measured by a quartz crystal thickness-meter (Sycon STM-100/MF) located in close proximity of the substrate holder. A much thicker film (250nm) was subsequently deposited on the other side of the etalon. The reflectivity of the deposited gold films were subsequently checked via both normal-incidence and off-normal incidence specular reflectance measurements using a rotating stage and visible optical spectrometer (QE-Pro, Ocean Optics), where they were confirmed to be 95 (+/- 1.5 )% and 100% respectively at the relevant wavelength.

*Radial Apodization filters:* The non-Gaussian intensity profile of light exiting a VIPA results in a non-optimal spectral projection. This can be compensated by employing a gradient Neutral Density (ND) filter, to effectively rectify the spatial intensity profile, resulting in an improved spectral projection and hence finesse. While for conventional VIPAs commercially available gradient ND filters may be employed [1], for the case of the radial VIPA a custom radial gradient ND-filter is required. To this end we fabricated custom radial gradient ND filters where the transmission is minimum in the center and drops off radially in an approximately Gaussian manner. Gradient-index gold coatings were deposited onto 4” borosilicate glass wafers by an in-house developed sputter unit equipped with a Kaufman-type argon ion source described above. The radial profile in the thickness of the Au was tuned by adjusting the distance between the center of the Au target (effectively acting as a point source) to the center of a rotating substrate holder. Doing so a radial Au thickness and hence transmission profile could readily be realized. The base pressure was 1.5 x $10^{-6}$ mbar, and the gold was deposited onto the surface at an average rate of 10 Å $s^{-1}$ at normal incidence. (The thickness and deposition rates of Au were subsequently monitored by quartz crystal thickness-meter directly facing the sputter source). Finally, the gradient-index Au films were annealed at a temperature of 773 K for 2 hours to smoothen the surface gradient profile. The masks were inserted into the setup at a slight angle to avoid back reflection. Though the radial profile is not optimum [1,2], it notably improved the spectral projection.

#### 2. Obtaining elastic moduli from $\nu_B$

The angular frequency $\omega$ of the probed phonons will be proportional to their phase velocity $V_{\boldsymbol{q}}$ in the direction of the scattering wavevector[3] $\boldsymbol{q}$:

$$\omega(\boldsymbol{q}) = qV_{\boldsymbol{q}} \qquad \textbf{Eqn. S1}$$

Where $q = |\boldsymbol{q}| = 2nk_0\cos(\theta/2)$, and $k_0 = 2\pi/\lambda_0$. Here $n$ is the refractive index of the sample, $\lambda_0$ is the free-space wavelength of the probing laser, and $\theta$ is the angle between the probing and detected direction. Writing the angular frequency in terms of the frequency of the phonons, which is also the frequency shift that is measured in BLS ($\omega = 2\pi\nu_B$), one can readily relate the measured BLS frequency shift to the hypersonic velocity:

$$\nu_B(\boldsymbol{q}) = \pm 2n\lambda_0^{-1}V_{\boldsymbol{q}}\cos(\theta/2) \qquad \textbf{Eqn. S2}$$

The propagation of the phonons can be described by a plane wave, for which the wave equation can be expressed in terms of the stiffness tensor $c_{ijkl}$:

$$\frac{\partial^2 u_i}{\partial t^2} = (c_{ijkl}/\rho)\ \frac{\partial}{\partial x_j}\left(\frac{\partial u_l}{\partial x_k}\right) \qquad \textbf{Eqn. S3}$$

where $\rho$ is the mass density, and $u$ is the local displacement in the indexed direction as a function of position $(x)$ and time $(t)$. The relevant stiffness tensor component will depend on the direction of propagation as well as whether the wave is a transverse or longitudinal wave. The coefficient $c_{ijkl}/\rho$ here is by design the square of the phonon velocity, which can be extracted via the Christoffel equation[3]:

$$\left|\hat{q}_k\hat{q}_l c_{ijkl} - \rho V_{\hat{\boldsymbol{q}}}^2\right| = 0 \qquad \textbf{Eqn. S4}$$

where $\hat{q}_k$ and $\hat{q}_l$ are the projections of the unit scattering wavevector ($\hat{\boldsymbol{q}} = \boldsymbol{q}/|\boldsymbol{q}|$) in the indexed direction. It follows that by measuring $\nu_B$ for different $\hat{\boldsymbol{q}}$ one can obtain the different components of $c_{ijkl}$, and thereby the mechanical anisotropy from Eqn.'s S2 and S4. The dominant contribution in polarization unresolved measurements comes from longitudinal modes, and measurements at different angles will typically probe combinations of the first three diagonal stiffness tensor components ($c_{iiii}$), as defined by the scattering wavevector, corresponding to the *longitudinal modulus* in the indexed directions $M_i = c_{iiii}$ for $i = 1,2,3$.

#### 3. Spatial, temporal and spectral resolution

*Spatial resolution:* The effective spatial resolution in BLS microscopy is in general non-trivial[4]. While the effective optical resolution can readily be calculated in the same way as with other optical confocal microscopy approaches (*i.e.* convolution of the excitation and detection), the resolution with which the acoustic velocities, and hence elastic moduli, can be measured may be dependent on the scales associated with the probed phonons. The highest possible resolution in the direction of the probed phonons can be estimated as $\sim\lambda_0/2n \approx 200$ nm, which one can interpret as the distance between two successive nodes/antinodes in the acoustic wave required to realize the necessary interference effect observed in BLS. In a spatially heterogeneous material with significant spatial variations in the acoustic impedance, this quickly becomes more complicated and reflections at acoustic boundaries as well as finite size effects/resonances may become relevant. Similarly, in materials with gradient properties (as are conceivable in biological structures), the plane wave approximation (Eqn. S3) breaks down and more elaborate molecular hydrodynamic explanation become necessary[5].

To obtain an estimate of the spatial resolution for our setup (Fig 1A, Fig S14) we look at the effect on the BLS frequency shift when scanning across a methanol – plastic interface, the results of which are presented in Fig S2. As can be seen the transition will be dependent on the probing angle relative to the interface, which can be explained in terms of the direction and portion of the probing volume occupied by the probed phonons (see also caption of Fig S2). The resolution in these tests measurements is $\approx 1.5\ \mu m$ , which can be assumed as a reasonable estimate of the resolution in the other measured samples.

The spatial resolution was in general found to be better in setup "A" than in setup "B" which was attributed to the angular misalignment of the excitation and detection point spread function resulting in a smaller effective probing volume, and to be expected.

The spatial resolution is important when measuring the properties of cell walls (*e.g.* Fig S2) to assure that the entire probed volume (effective point spread function) can be associated with the cell walls. It is also for this reason that setup "B" was employed for measuring the periclinal walls (despite the overall lower resolution compared to setup "A") since both the excitation and detection PSF where at very large angles and thus can be considered not to extend significantly into the surrounding cell cytoplasm.

The spatial resolution also becomes critical when considering spatial correlations (such as in Fig 3 and Fig 4). A reduced spatial resolution can be expected to result in increased small-distance correlations, and as such limits how fine one can perform grid scans. In terms of the interpretation of results, a reduced resolution would accentuate an exponential distance scaling rather than an algebraic scaling. For this reason grid steps exceeding the spatial resolution where employed.

The spatial resolution with which the linewidth can be measured depends on the attenuation distance of the probed phonons (typically 1-2$\mu m$)[6]. In our studies we only study the linewidth of periclinal walls (which were significantly larger than these distances in the probed directions, i.e. almost perpendicular to the optical axis). We furthermore did not perform any spatial correlation analysis and as such the precise spatial resolution would not be significant in these cases. Finally, multiple scattering can also affect the linewidth significantly in scattering samples[7]. However since we are always measuring close to the surface these can be assumed to be small.

*Temporal resolution:* In practice numerous factors will affect the temporal resolution, including the probing laser power, the BLS scattering cross-section for that particular sample region, and the accuracy with which one wishes to localize the BLS peaks or determine the peak-widths. As such a compromise has to be reached between how accurately one wishes to measure the spectra and how much (laser) energy one can apply without inducing non-physiological perturbations that affect the measurement or studied processes, which will be sample and problem specific. The laser power at the sample position in our case was in all cases limited to 10 (+/-2) mW or lower as described in *Material & Methods*. The acquisition time per position was varied from <100ms to several seconds depending on the setup, as indicated for each individual sample. These are comparable to those employed in standard Brillouin microscopy experiments on isolated mammalian cells [1] and live *Arabidopsis* hypocotyl plant cells [8] where they were confirmed to not induce noticeable photo-toxicity over measurement time scales. It should also be noted that due to the nature of the performed measurements (point measurements on comparatively coarse grid scans) the total energy deposited on the sample in our measurements is significantly (often orders of magnitude) less than in the quoted studies where much larger and finer-grid point scanning was used to render high resolution spatial maps.

*Spectral resolution:* The spectral resolution of VIPA-based Brillouin spectrometers will in general be worse than scanning multi-pass Fabry-Perot based spectrometers[6]. Deconvolution with the instrument spectral response (see *Materials & Methods*) however significantly improves the spectral resolution, as will the Apodization filtering[1] and Fourier space filtering[9] - both employed in our setup (Fig S14). The accuracy with which the Brillouin frequency shift $\nu_B$ can be obtained will ultimately be determined by the accuracy with which one can localize the BLS peak position (in a comparable manner to that in superresolution fluorescence localization microscopy for localizing the position of single emitters)[10]. To this end increased photon statistics (as can be realized using longer acquisition times and higher laser powers) can result in a higher spectral precision. In our case the uncertainty associated with the fitting is significantly smaller than the sample-sample statistical variability observed. To this end all error bars presented, when visible and larger than the plotted data point, are determined essentially by the statistical uncertainty from repeated measurements unless indicated otherwise.

The issue of spectral resolution becomes more acute when one wishes to also determine the peak width (as is the case for the loss tangent in *e.g.* Fig 2). To this end extra care needs to be taken to deconvolve data with the spectral response not just of the instrument but also of the instrument-sample combination (*i.e.* complete optical path). This is done by measuring the elastic scattered light at the same sample position and using this for the deconvolution kernel (with the assumption that this is the system response to a true single-frequency delta-function at the probing wavelength) (Methods). Given the spectral width of the probing laser is < MHz, and the expected small contribution due to other scattering processes (*e.g.* Rayleigh peak, Mountain peak), this can be assumed a reasonable assumption.

#### 4. Effective Medium modeling

*Muscle myofibers:* The anisotropy data for the Brillouin frequency shift of the myofibers was analyzed by considering the projection of the scattering wavevector on the stiffness tensor, with the function:

$$\nu_B(\phi) = \rho^{-1/2} \left(\sum_{i=1,2,3} q_i^2 c_{ii}\right)^{1/2} \quad \textbf{Eqn. S5A}$$

For setup "B" (Methods) $q_i$ are given by:

$$\begin{bmatrix} q_1 \\ q_2 \\ q_3 \end{bmatrix} = \frac{2n}{\lambda_0} \begin{bmatrix} \sin(\phi - \phi_0)\sin(\theta_s) \\ \cos(\phi - \phi_0)\sin(\theta_s) \\ \cos(\theta_s) \end{bmatrix} \quad \textbf{Eqn. S5B}$$

Where $n$ and $\rho$ are the refractive index and mass density, and $\phi$ is the azimuthal angle. $\phi_0$ is an azimuthal off-set angle (such that the coordinate axis aligns with the transverse anisotropic symmetry axis, *i.e.* fiber orientation) which is assigned from the orientation of the fiber determined from the widefield image. $\theta_s$ is the (fixed) polar probing angle specific to the setup and objective lens used. For setup "A" (Methods), $\theta_s$ would be replaced by $\theta_s/2$. $\theta_s$ is determined from the distance of the hole in the rotating disk to the center of the disk in setup "B", or the radius of the annular mask *r* in setup "A" (see Fig S1A and Fig S1B) relative to the diameter of the back aperture of the objective lens (*d*). Given the numerical aperture (*NA*) of the objective lens, and the sample refractive index *n* (typically between 1.8-1.4), this can be obtained via: $\theta_s \approx \sin^{-1}(2r \cdot d^{-1} \cdot n^{-1} \cdot NA)$. For the case of the myofibers the transverse anisotropic mechanical properties would imply $c_{22} = c_{33}$. It follows from Eqns. S5 that the values of $c_{11}, c_{22}$ and $c_{33}$ are obtained by fitting $[\nu_B(\phi)]^2$ with:

$$[\nu_B(\phi)]^2 = A[B_1 \sin^2(\phi - \phi_0)\, c_{11} + [B_1 \cos^2(\phi - \phi_0) + B_2] c_{22}] \quad \textbf{Eqn. S6}$$

for $c_{11}$ and $c_{22}$. $A = 4n^2\lambda_0^{-1}\rho^{-1}$, $B_1 = \sin^2(\theta_s)$, and $B_2 = \cos^2(\theta_s)$ are fixed numerical constants.

To consider the hydration dynamics we consider the fiber consists of a solid and liquid portion, with the former structured in a manner that the effective material response will be either iso-stress (parallel to fiber) or iso-strain (perpendicular to fiber). From Eqn. S6, one may write $M_V = c_{11}$ and $M_R = c_{22}$ parallel and perpendicular to the fiber axis respectively, where $M_V$ and $M_R$ are the Voigt and Reuss effect responses (see Methods). Associating $M_1 = M_s$ with the solid component ($M_s$) and $M_2 = M_L$ with the liquid component ($M_L$), it follows that:

$$M_V = M_L + M_s(1 - M_L M_R^{-1})$$

This implies the scaling of the two stiffness tensor components should follow $c_{11}^{-1} \propto c_{22}$, with a slope given by $\frac{dM_R^{-1}}{dM_V} = -(M_s M_L)^{-1}$ if the change in Moduli is exclusively due to changes in the liquid filling fraction. This appears to be the case, as is shown Fig S3C where the two parameters are plotted against each other, suggesting that the transverse anisotropic symmetry is maintained during dehydration. The values proportional to $c_{11}$ and $c_{22}$ here were obtained from fitting Eqn. S6 to $\nu_B(\phi)$ (Supplementary Fig 3B). We note that here we have assumed the refractive index to be isotropic (which is almost certainly not the case) and the density to be constant. Under these assumptions we may using the above attempt to extract $M_s$ and $M_L$. For the latter we obtain $M_L \approx 2.5\ GPa$, which can be considered reasonable, for the solid component ($M_S$) we obtain a value almost an order of magnitude larger, which is unlikely, and could be due to the assumptions of a constant isotropic refractive index and constant density during dehydration. Indeed, assuming the refractive index is larger parallel to the fiber axis will significantly reduce $M_S$. However, given that the degree to which this is the case is still unknown.

From the above may also readily write:

$$\frac{M_V}{M_R} = 1 + \bar{M}_s^{-1}(\bar{M}_s - 1)^2(1-f)f \approx \left(\nu_B^{max}/\nu_B^{min}\right)^2 \left(n_\perp/n_{||}\right)^2,$$

where $\bar{M}_s = M_S/M_L$. As a first approximation during dehydration the solid fraction $f$ can be assumed to scale as $\propto (1 - e^{-(t+\Delta)/\tau})$ where $t$ is the time, $\Delta$ is a temporal offset, and $\tau^{-1}$ is the effective dehydration rate. The dehydration data (Supplementary Fig 3A) is well fit with this equation with the constraints $\{\bar{M}_s, \Delta, \tau > 0\}$ and $\{0.5 \leq n_\perp/n_{||} \leq 2\}$.

The sparseness of our knowledge for the mechanical properties and relaxation processes of cell walls in the probed GHz-frequency regime in Arabidopsis cell walls, make rigorous quantitative modeling of the BLS mechanical anisotropy challenging. The semi-quantitative analysis pursued here thus hinges on known structural features and (a)symmetries, which are assumed to be reflected to some degree in the BLS measured properties, together with phenomenological parameters that describe the local and molecular scale interactions (*e.g.* cross-linking) expected to contribute to the BLS measured properties. A means of describing measured BLS moduli in compound materials involves the use of effective medium approximations, which have been used to both account for the effect of sub-wavelength structural features as well as the nature of interactions between constituent molecules[11,12]. Given the small dimensions (<<100 nm) of constituent anisotropic structures such as microfibrils in the cell wall, such an approach can be justified for BLS measurements.

The cellulose microfibrils in the cell wall are considered to be the load-bearing structure that define the anisotropic mechanics, although there is active debate regarding their role in defining cell morphology (see main text). Nevertheless, their distinct orientational distributions in *e.g.* periclinal and anticlinal walls, while not always as pronounced in epidermal cells (probed in our studies), present a good starting point for modelling the anisotropic mechanical properties. To this end, a Voigt and Reuss model (Methods) describing the effective mechanical response when the average fibril alignment is perpendicular or parallel to the scattering wavevector serves as the simplest starting point. Deviations from these "ideal" models would however be expected (see below).

For the Voigt and Reuss models we employ Eqn. S6 where $M_1 = M_C$ and $M_2 = M_M$ are the longitudinal moduli for the cellulose fibrils and the surrounding matrix respectively, and $f$ is the microfibril cellulose fraction. It is reasonable also to assumed that $M_C \gg M_M$. Noting that $f \approx 0.7$ [13], it is straight away clear (from Eqn S6) this would result in $M_V \gg M_R$ and thus an unrealistically high anisotropy--i.e. $\nu_B^{(\times)}(\phi_0) \gg \nu_B^{(\times)}(\phi_0 + \pi/2)$, that is not observed in our BLS measurements (Fig 2). There are several potential reasons for this discrepancy. Firstly, it is possible the contrast between the BLS-moduli $M_C$ and $M_M$ is not as stark as it would be in the quasi-static tensile regime. This is possible given the effective solid like properties soft matter takes on at high frequencies. Secondly, the microfibrils are in practice not perfectly aligned (especially in the epidermal cells), and there will exist a distribution of microfibril angles as well as potentially a contribution from random microfibril orientations. To a first approximation we can account for the latter by including an additional isotropic term $c_{ii} \rightarrow c_{ii} + M_0$ , with the constraint that $M_0 < (M_R + M_V)/2$ (see below). Thirdly, the simple effective medium model employed does not take into account the effects of cross-linking of the microfibrils (*e.g.* from xyloglucan hemicellulose), which can be expected to be significant in the BLS regime. The effect of cross-linking can be expected to be most significant perpendicular to the microfibril axis, for which a correction may be applied by replacing $M_R \rightarrow (M_R^{-1} + M_{CL}^{-1})^{-1}$ where $M_{CL}$ is the contribution to the modulus from cross-linking. While one can expect shear moduli in the direction parallel to the microfibril axis to be affected by cross-linking, the effective longitudinal moduli, $M_V$ would be affected to a lesser degree. Including this contribution, together with the contribution from random orientated microfibrils, we may write the frequency shift for the outer ("×" in Supplementary Fig 4A) and inner ("+"in Supplementary Fig 4A) walls, in the coordinate system illustrated in Supplementary Fig 4A:

$$\begin{pmatrix} [\nu_B^{(\times)}(\phi)]^2 \\ [\nu_B^{(+)}(\phi)]^2 \end{pmatrix} = \rho^{-1} \begin{pmatrix} M'_V q_1^2(\phi) + M'_R[q_2^2(\phi) + q_3^2(\phi)] \\ M'_V q_3^2(\phi) + M'_R[q_1^2(\phi) + q_2^2(\phi)] \end{pmatrix} \qquad \textbf{Eqn. S7A}$$

$$M'_V = (1-\alpha)M_V + \alpha M_0 \quad \text{and} \quad M'_R = (1-\alpha)(M_R + M_{CL}) + \alpha M_0 \qquad \textbf{Eqn. S7B}$$

where $0 < \alpha < 1$ is the isotropic contribution from randomly aligned fibers, and $q_i$ are as defined in Eqn. S5B, and can readily identify the stiffness tensor components as (see Supplementary Fig 4A):

$$c_{11} = M'_V \text{ and } c_{22} = c_{33} = M_R{}', \qquad \textbf{Eqn. 8A}$$

$$c_{11} = c_{22} = M'_R \text{ and } c_{33} = M'_V. \qquad \textbf{Eqn. 8B}$$

It follows from Eqn. S7A that:

$$[\nu_B^{(\times)}(\phi)]^2 = A[B_1 \sin^2(\phi - \phi_0) M'_V + [B_1 \cos^2(\phi - \phi_0) + B_2]]M'_R \qquad \textbf{Eqn. 9A}$$

$$[\nu_B^{(+)}(\phi)]^2 = A(B_2 M'_V + B_1 M'_R) \qquad \textbf{Eqn. 9B}$$

where $A = 1.378 \cdot 10^{13}\ kg/m$, $B_1 = 0.2204$, $B_2 = 0.7796$, and we have taken the values of $\rho = 1100\ kg/m^3$ [14] and $n = 1.41$ [15].

The outer walls can then be fitted with Eqn. 9A for $M'_V$, $M'_R(< M'_V)$ and $\phi_0$ (solid line in Supplementary Fig 4B). They may also be fitted only with $M'_V$, and $\phi_0$ with the constraint from Eqn. 9B, *i.e* $M'_R = B_1^{-1}(A^{-1}[\nu_B^{(+)}]^2 - B_2 M'_V)$ where $\nu_B^{(+)}(\phi) = \langle \nu_B^{(+)}(\phi) \rangle_\phi = \nu_B^{(+)}$ is the angular average of the BLS frequency shift from the inner walls (dashed line in Supplementary Fig 4B). It is clear that the former fit is much better, and that the constraint from the inner wall vastly overestimates $M'_R$ for the outer wall. This suggests that $M'_V/M'_R$ is *smaller* for the outer wall, namely that it is less anisotropic. The differences in the obtained $M'_V/M'_R$ are shown in Supplementary Fig 4C.

This could be the result of: ***1.*** *Change in Microfibril density:* There is the possibility of a different microfibril density ($f$) between the two cell wall types studied. This could occur because the stress experienced by the outer wall ("×" in Fig 2A) is distinct from that of the inner wall ("+" in Fig 2A), resulting in a different strain and in turn different microfibril density [16]. A higher in-plane strain in the outer wall may be expected to result in an effective decrease in fiber density ($f$), and lead to an increase of $M_R$ relative to $M_V$ as observed. ***2.*** *Distinct Microfibril angular distribution:* A different orientation and/or a distinct angular distribution of orientations for the microfibrils between the walls cannot be ruled out. This is however less likely to be significant, given that high resolution 3D confocal fluorescence studies, as well as small-angle X-ray scattering studies, which indiscriminately probe the orientations of microfibrils in both studied cell wall orientations, suggest a common orientation axis [17]. ***3.*** *Distinct cross-linking:* A different degree of cross-linking in the outer walls versus the inner walls can be expected to result in a distinct $M_V' / M_R'$. ***4.*** *Refractive index anisotropy:* Based on the subwavelength structural asymmetry, the refractive index can be expected to have a corresponding anisotropy. This may be explicitly considered by replacing $q_i \rightarrow n_i n_0^{-1} q_i$, where $n_i$ is the refractive index in the indexed direction and $n_0$ is the angle averaged value. While a difference in this anisotropy (which would be a consequence of ***1-3***) will in itself also result in a difference between $M_R'$ and $M_V'$ , there is currently no data for this, and it can thus not be ruled out.

#### 5. Association of Reuss vs. Voigt scaling with ionic concentration

The Reuss and Voigt models (described above for calculating effective mechanical properties in transverse isotropic media), can also give qualitative insight into the nature of hydrostatic interactions/screening [12,18] and their effect on the BLS measured properties. For the case of negatively charged polymers in a solution, when the solution has a low cationic concentration, the polymers will couple more effectively to the solvent, and the system can be expected to scale more like a Voigt-effective medium with increasing cation concentration. On the other hand, for high cationic concentrations the screening from cations will reduce the effectiveness with which polymers couple to both the solvent and each other. As a result, the system can be expected to scale more like a Reuss effective medium with increasing cation concentration. If the cation concentration is assumed to be related to the amount of chromatin condensation, it follows that as a function of chromatin condensation (which scales with the SIR-DNA fluorescence intensity on a single cell level, see Supplementary Fig 7C), one would for a low cationic concentration (more decondensed chromatin) expect scaling to more closely resemble a Voigt model. On the other hand, for a higher cation concentration (more condensed chromatin) one may expect a Reuss-like scaling. To approximate this behavior, in Supplementary Fig 9 a weighted model is assumed (Eqn. 7), with the weighting factor directly equal to the normalized fluorescence intensity (*i.e.* condensed chromatin fraction). This will describe a Voigt-like scaling for low condensed chromatin and Reuss-like scaling for high condensed chromatin fractions. This results in an increasing and then saturating and slightly decreasing dependence of the modulus with increasing condensed chromatin fraction (*viz.* cationic concentration). While the true functional form is likely not this trivial, the general trend of increasing and then saturating effective modulus with increasing fluorescence intensity will result also for deviations from this simple proportionality. Also shown in Supplementary Fig 9 are the case of a pure Voigt and a pure Reuss scaling, which would predict a continuous linear and polynomial increase in the modulus.

Our data shows an increase and then saturated (followed by a possible slight decrease) of the modulus with increasing chromatin condensation (Supplementary Fig 9), consistent with the low and high chromatin condensation fraction corresponding to low and high cation concentrations respectively. This is consistent with the proposed model (main text) in that local changes in cation concentration are correlated to the condensed chromatin fraction which are in turn related to the BLS in living cell nuclei. It is worth noting is that the correlation between increased (steady state) fluorescence intensity for increased condensed chromatin is only true on a cell per cell basis (as shown in Supplementary Fig 7, see also[19]). The analysis in Supplementary Fig 9 was thus performed by normalizing the fluorescence intensity for each cell separately. Not normalizing the fluorescence intensity (or normalizing this after averaging over cells), revealed no significant correlation between the SIR-DNA fluorescence intensity and the mean Brillouin frequency shift. This can be taken to suggest that the SIR-DNA fluorescence intensity is only a good indicator of the fraction of condensed chromatin on a single cell level (consistent with conclusions drawn from Supplementary Fig 7A & 7C).

#### 6. 3-species reaction diffusion model

A simple heuristic three-species model, shown in Fig 4C, of condensed chromatin (C), positive ions (+), and a larger diffusive species which inhibits chromatin condensation and serves as a crowing agent (X), can support stable Spatial and Temporally Oscillating Solutions (STOS). While it is possible to construct a two species reaction-diffusion model (consisting of *e.g.* only cations and condensed chromatin) that exhibits STOS, this would require non-linear interaction terms and additional assumptions that cannot be as easily motivated.

The core assumptions of this rather simple model are that increasing the cation concentration will increase chromatin condensation, the effectiveness of which is modulated by a mobile inert macromolecular species/crowding agent (that is at this point still undefined). While the effect of an inert crowding agent may include volume seclusion effects, reducing diffusion rates, and enhancement of binding rates[20], here we only consider its effect on the effective interaction volume. The specific cationic species is also left undefined (in practice different species will have differing effects on chromatin condensation[21]). The employed model is a simplification also insofar that it includes no consideration of the detailed structural organization of chromatin. It further assumes unconstrained diffusion of cations and the crowding agent (which is also not the case, although on longer time scales it is a reasonable assumption[22]). It assumes single rate constants for each of the processes (in practice one would expect this to be a spectrum due to multiple species and the heterogeneous environment), and makes no allusions as to whether these are in of themselves passive or active processes. It nevertheless may serve useful for understanding how such species affect spatial and temporal scaling of the chromatin state, and as we find, can result in STOS that can be related to those observed in our experiments. Below we describe the key assumptions of this model and motivation for the different reaction pathways.

Since the effects of crowding are not implicitly accounted for, the cation concentration refers to the effective cation concentration, namely that calculated by considering the volume that is available for it to occupy (which is decreased by an increase in the crowding agent). As such an increase in crowding agent concentration increases the effective cation concentration.

Chromatin diffusion coefficients are 3-4 orders of magnitude smaller than any proteins in the nucleus [23]. In our model the condensed chromatin is thus assumed to be effectively immobile relative to the cations and the crowding agent, which are assumed to be diffusive (as indicated by squiggly lines in Fig 4C).

An increase in the concentration of cations increases the condensed chromatin concentration (activation arrow between the cation and the chromatin in the graph) – this is an extensively documented effect, although again it will be more complex and depend on cation species[21,24].

When chromatin condenses the effective volume available to the cations and the crowding agent is increased, such that their effective concentration is reduced (inhibition arrows from the chromatin to the cations and crowding agent). This inhibitory affect may be considered weak and smaller than the cation/condensed-chromatin activation.

The effect of the crowding agent in relation to chromatin condensation and effective ion concentration is non-trivial. The former especially is experimentally challenging to disentangle from a changing effective cation concentration. On the one hand it can be expected to have an inhibitory effect on chromatin condensation by virtue of volume exclusion effects that inhibit the effective screening of cations, which would cause condensation [25,26] . On the other hand, it has also been proposed to inhibit intra-chromosome interactions (restoring the 10nm chromatin fiber structure and mimicking an increasing cation concentration), suggesting the opposite effect [27]. The former and latter can be understood as being dependent on the size of crowding agent. Here we assume a weak inhibitory effect (inhibitory arrow between crowding agent and the condensed chromatin) that is much smaller than other inhibition rates, since in the opposite case we cannot find STOS.

In relation to the cation concentration, an increase in the crowding agent is assumed to result in an increase in the effective cation concentration due to the crowding agent reducing the effective volume that can be occupied by cations (activation arrow between crowding agent and cations). This activation is considered to be smaller than the cation/condensed-chromatin activation but larger than the inhibition effects of condensed chromatin on the cations and crowding agent.

An increase in the crowding agent or the cation concentration will in themselves limit their respective subsequent increase, due to electrostatic, volumetric, and/or osmotic effects (inhibition loops for cations and for crowding agent). These are both considered smaller than the cation/chromatin and crowding-agent/cation activation.

Since the total concentration of chromatin can be assumed constant, a weak condensed-chromatin self-inhibitory effect was included to account for the fact that we are describing the condensed chromatin state concentration (which cannot exceed the total local chromatin concentration since the chromatin is assumed to be effectively immobile) and which will effectively be depleted as more chromatin changes from decondensed to condensed.

The diffusion coefficient of the cations was set to be larger than the crowding agent based on the assumption that the latter is larger in size.

It is important to note that in our model the concentration of cations and crowding agent represent the effective concentration in the available interaction volume. The condensed chromatin condensation on the other hand will be related to the fraction of chromatin in a "condensed state". Since it is taken to be immobile, a local increase (decrease) thereof would thus correspond to chromatin changing from a more decondensed (condensed) to a condensed (decondensed) state.

Writing the activation rate of species $i$ by species $j$ as $k_{ij}$, and the diffusion coefficient of species $i$ as $D_i$, where " $C$ " =condensed chromatin, " + " =cations, and " $X$ " =crowding agent, we find STOS can exist when: $\{D_C \approx 0\} \ll D_X \approx 0.1\ D_+$ ; $k_{XC} \approx 0.1 k_{C+}$ ; and $k_{++} \approx 3|k_{XX}|$. We further require: $k_{CX} < 0$; $k_{C+} > k_{+X} > 0$; $\{|k_{CC}|, |k_{XC}|, |k_{+C}|\} < k_{+X}$; $|k_{++}| < |k_{CC}|$ and $\{ k_{++}, k_{XX}, k_{CC}, k_{XC}, k_{+C} \} < k_{CX}$. Given $k_{CX} < 0$ the crowding agent(s) inhibit chromatin condensation which together with their "fast" diffusion $D_X \approx 0.1\ D_+$ is consistent with them being (predominantly) relatively small molecules. The condition $k_{++} \approx 3|k_{XX}|$ also suggests their self-inhibition is significant. Unlike cations (where electrostatic interactions may justify a significant passive self-inhibition) this is less likely to be the case for the crowding agent(s), and suggests an active mechanism for their regulation.

Taking the diffusion of cations in the nucleus as $D_+ \sim 500 \mu m^2 s^{-1}$ this would imply a diffusion coefficient for the crowding agent of $D_X \sim 50 \mu m^2 s^{-1}$. One may use this to roughly estimate the molecular size[28] as $\sim$ 50kDa. There are numerous candidates for proteins known to cause decondensation of this size, including ATPases such as RuvBL 1/2 (50kDa) [29], p97 (90kPa) [30], and various Hystone Acetylasetrasferases (HATs) (40-100s kDA)[31]. Decondensation of chromatin by all these is either in of itself an active process (requiring ATP and GTP hydrolysis) and/or is actively regulated, which may provide the required mechanism (and energy) for maintaining critical behavior and STOS. Indeed, the disruption of regulatory processes associated with these proteins are known to drastically affect homeostasis.

#### 7. Supplementary References


1 Scarcelli, G. *et al.* Noncontact three-dimensional mapping of intracellular hydromechanical properties by Brillouin microscopy. *Nat Methods* **12**, 1132-1134 (2015). https://doi.org:10.1038/nmeth.3616

2 Antonacci, G., De Panfilis, S., Di Domenico, G., DelRe, E. & Ruocco, G. Breaking the Contrast Limit in Single-Pass Fabry-P\'erot Spectrometers. *Physical Review Applied* **6**, 054020 (2016). https://doi.org:10.1103/PhysRevApplied.6.054020

3 Berne, B. J. & Pecora, R. *Dynamic Light Scattering: With Applications to Chemistry, Biology, and Physics*. (Dover Publications, 2000).

4 Caponi, S., Fioretto, D. & Mattarelli, M. On the actual spatial resolution of Brillouin Imaging. *Opt Lett* **45**, 1063-1066 (2020). https://doi.org:10.1364/OL.385072

5 Boon, J. P. & Yip, S. *Molecular Hydrodynamics*. (Dover Publications, 1991).

6 Antonacci, G. *et al.* Recent progress and current opinions in Brillouin microscopy for life science applications. *Biophysical Reviews* (2020). https://doi.org:10.1007/s12551-020-00701-9

7 Mattarelli, M., Capponi, G., Passeri, A. A., Fioretto, D. & Caponi, S. Disentanglement of Multiple Scattering Contribution in Brillouin Microscopy. *ACS Photonics* **9**, 2087-2091 (2022). https://doi.org:10.1021/acsphotonics.2c00322

8 Elsayad, K. *et al.* Mapping the subcellular mechanical properties of live cells in tissues with fluorescence emission-Brillouin imaging. *Sci Signal* **9**, rs5 (2016). https://doi.org:10.1126/scisignal.aaf6326

9 Edrei, E., Gather, M. C. & Scarcelli, G. Integration of spectral coronagraphy within VIPA-based spectrometers for high extinction Brillouin imaging. *Opt Express* **25**, 6895-6903 (2017). https://doi.org:10.1364/OE.25.006895

10 Török, P. & Foreman, M. R. Precision and informational limits in inelastic optical spectroscopy. *Scientific Reports* **9**, 6140 (2019). https://doi.org:10.1038/s41598-019-42619-7

11 Wu, P. J. *et al.* Water content, not stiffness, dominates Brillouin spectroscopy measurements in hydrated materials. *Nat Methods* **15**, 561-562 (2018). https://doi.org:10.1038/s41592-018-0076-1

12 Bailey, M. *et al.* Viscoelastic properties of biopolymer hydrogels determined by Brillouin spectroscopy: A probe of tissue micromechanics. *Science Advances* **6**, eabc1937 (2020). https://doi.org:10.1126/sciadv.abc1937
13 Thompson, D. S. How do cell walls regulate plant growth? *Journal of Experimental Botany* **56**, 2275-2285 (2005). https://doi.org:10.1093/jxb/eri247
14 Rapusas, R. S. & Driscoll, R. H. Thermophysical properties of fresh and dried white onion slices. *Journal of Food Engineering* **24**, 149-164 (1995). https://doi.org:https://doi.org/10.1016/0260-8774(94)P2640-Q
15 Liu, D. Y. *et al.* Reflection across plant cell boundaries in confocal laser scanning microscopy. *J Microsc* **231**, 349-357 (2008). https://doi.org:10.1111/j.1365-2818.2008.02068.x
16 Cosgrove, D. J. Diffuse Growth of Plant Cell Walls. *Plant Physiology* **176**, 16-27 (2018). https://doi.org:10.1104/pp.17.01541
17 Saxe, F. *et al.* Measuring the distribution of cellulose microfibril angles in primary cell walls by small angle X-ray scattering. *Plant Methods* **10**, 25 (2014). https://doi.org:10.1186/1746-4811-10-25
18 Adichtchev, S. V. *et al.* Brillouin spectroscopy of biorelevant fluids in relation to viscosity and solute concentration. *Phys Rev E* **99**, 062410 (2019). https://doi.org:10.1103/PhysRevE.99.062410
19 Hockings, C. *et al.* Illuminating chromatin compaction in live cells and fixed tissues using SiR-DNA fluorescence lifetime. *bioRxiv*, 2020.2005.2002.073536 (2020). https://doi.org:10.1101/2020.05.02.073536
20 Hancock, R. in *International Review of Cell and Molecular Biology* Vol. 307 (eds Ronald Hancock & Kwang W. Jeon) 15-26 (Academic Press, 2014).
21 Strick , R., Strissel , P. L., Gavrilov , K. & Levi-Setti , R. Cation–chromatin binding as shown by ion microscopy is essential for the structural integrity of chromosomes. *Journal of Cell Biology* **155**, 899-910 (2001). https://doi.org:10.1083/jcb.200105026
22 Höfling, F. & Franosch, T. Anomalous transport in the crowded world of biological cells. *Reports on Progress in Physics* **76**, 046602 (2013). https://doi.org:10.1088/0034-4885/76/4/046602
23 Bornfleth, H., Edelmann, P., Zink, D., Cremer, T. & Cremer, C. Quantitative motion analysis of subchromosomal foci in living cells using four-dimensional microscopy. *Biophys J* **77**, 2871-2886 (1999). https://doi.org:10.1016/S0006-3495(99)77119-5
24 Sen, D. & Crothers, D. M. Condensation of chromatin: role of multivalent cations. *Biochemistry* **25**, 1495-1503 (1986). https://doi.org:10.1021/bi00355a004
25 Yu, T., Zhu, Y., He, Z. & Chen, S.-J. Predicting Molecular Crowding Effects in Ion–RNA Interactions. *The Journal of Physical Chemistry B* **120**, 8837-8844 (2016). https://doi.org:10.1021/acs.jpcb.6b05625
26 Zimmerman, S. B. & Harrison, B. Macromolecular crowding increases binding of DNA polymerase to DNA: an adaptive effect. *Proceedings of the National Academy of Sciences* **84**, 1871-1875 (1987). https://doi.org:doi:10.1073/pnas.84.7.1871
27 Maeshima, K., Tamura, S., Hansen, J. C. & Itoh, Y. Fluid-like chromatin: Toward understanding the real chromatin organization present in the cell. *Current Opinion in Cell Biology* **64**, 77-89 (2020). https://doi.org:https://doi.org/10.1016/j.ceb.2020.02.016
28 Dross, N. *et al.* Mapping eGFP Oligomer Mobility in Living Cell Nuclei. *PLOS ONE* **4**, e5041 (2009). https://doi.org:10.1371/journal.pone.0005041
29 Magalska, A. *et al.* RuvB-like ATPases Function in Chromatin Decondensation at the End of Mitosis. *Developmental Cell* **31**, 305-318 (2014). https://doi.org:https://doi.org/10.1016/j.devcel.2014.09.001
30 Mérai, Z. *et al.* The AAA-ATPase molecular chaperone Cdc48/p97 disassembles sumoylated centromeres, decondenses heterochromatin, and activates ribosomal RNA genes. *Proceedings of the National Academy of Sciences* **111**, 16166-16171 (2014). https://doi.org:10.1073/pnas.1418564111

31 Ruan, K. *et al.* Histone H4 acetylation required for chromatin decompaction during DNA replication. *Scientific Reports* **5**, 12720 (2015). https://doi.org:10.1038/srep12720
